# Multi-omic analysis reveals maturation programs in human pluripotent stem cell-derived cardiomyocytes during long-term culture

**DOI:** 10.64898/2026.06.26.734802

**Authors:** Austin K. Feeney, Aaron D. Simmons, Elizabeth F. Bayne, Yanlong Zhu, Chanhyung Park, Claire J. Peplinski, Fathima Shabnam, Xiaotian Zhang, Jianhua Zhang, Melissa R. Pergande, Timothy J. Kamp, Ying Ge, Sean P. Palecek

## Abstract

Human pluripotent stem cell-derived cardiomyocytes (hPSC-CMs) hold tremendous promise for disease modeling, drug discovery, and cardiac regenerative therapies. However, the immature phenotype of hPSC-CMs remains a major barrier limiting their translational utility. Here, we performed integrated multi-omic profiling to identify molecular pathways and regulatory programs associated with hPSC-CM maturation during long-term culture. hPSC-CMs were cultured for 113 days and analyzed using metabolomics, proteomics, and transcriptomics across progressive stages of maturation. Long-term culture induced widespread multi-omic remodeling, including significant changes in 142/934 metabolites, 550/3,556 proteins, and 2,892/23,309 transcripts from Day 30 to Day 113. Metabolomic analyses revealed early increases in phospholipid biosynthesis and mitochondrial beta oxidation of fatty acids from Day 30 to Day 60, suggesting metabolic priming precedes later maturation events. In contrast, proteomic remodeling was more prominent during later stages of maturation and was characterized by enhanced calcium handling and cell cycle exit. Transcriptomic analyses demonstrated progressive increases in ion channel expression, t-tubule organization, fatty acid metabolism, creatine shuttle pathways, and cell cycle arrest programs. Transcriptomic and integrative multi-omic pathway analyses identified coordinated suppression of TGFβ, MAPK, Wnt, and Hedgehog signaling together with activation of integrin-related, respiratory electron transport, muscle contraction, and Slit-Robo signaling pathways during maturation. Moreover, multi-omic transcription factor activity analysis prioritized a GATA4-centered network of putative cardiomyocyte maturation regulators including SOX7, SOX18, TBX2, and ZFPM2 (FOG2). Together, these findings elucidate the degree and pace of hPSC-CM maturation during long-term culture and establish an integrated multi-omic framework for identifying strategies to accelerate hPSC-CM maturation.

## 1. Introduction

Human pluripotent stem cell-derived cardiomyocytes (hPSC-CMs) display considerable potential for modeling cardiac diseases^1,2^, developing pharmaceutical candidates^3,4^, and regenerating the heart^5^, but their utility is plagued by an immature phenotype compared to adult cardiomyocytes. Although hPSC-CM platforms have been widely adopted as an *in vitro* model since the emergence of hPSC-CM differentiation protocols^6–8^, there have been very few FDA approvals for novel cardiac drugs based on these models^9^. Notably, hPSC-CM maturation status affects drug and hypoxia sensitivity, underscoring the importance for maturation considerations in hPSC-CM drug development and disease modeling applications^2,10^. Moreover, clinical hPSC-CM applications continue to be hampered by safety concerns related to transplantation-induced arrhythmias arising from CM heterogeneity and immaturity^11,12^. Consequently, understanding and advancing CM maturation remains of paramount importance for evolving hPSC-CMs towards relevant models for improving the treatment of human heart disease.

The temporal transition to a mature CM phenotype requires distinct shifts in structural, electrophysiological, and metabolic cellular machinery to meet the complex demands of postnatal myocardium^13,14^. Structural maturation hallmarks include increased cell size, transition to a characteristic rod-like morphology, and increased sarcomere length and organization^15^. Sarcomeric maturation is also marked by characteristic isoform switches in the expression of fetal isoforms to adult isoforms such as the transition from MYL7 to MYL2 and MYH6 to MYH7^16,17^. Hallmarks of electrophysiologic maturation include refined calcium handling kinetics and specialized ion channel expression, which collectively support synchronized excitation-contraction coupling^18^. During CM maturation, metabolism shifts from reliance on glycolysis to more efficient energy sources including fatty acid oxidation and oxidative phosphorylation to sustain the high energy requirements of the heart^19^. Because fatty acid oxidation and oxidative phosphorylation generate ATP in mitochondria, this process necessarily corresponds with increased mitochondrial biogenesis and ATP transfer through the creatine shuttle pathway^20^.

Approaches to direct hPSC-CM maturation have focused on recapitulating aspects of the myocardial niche through micropatterned substrates to promote structural alignment^21,22^, electromechanical stimulation to enhance synchronized force generation^22,23^, 3-dimensional multi-cellular tissues to replicate native cellular architecture^24–26^, and media modifications to support metabolic and ionic requirements^16,22,27^. However, a majority of these strategies are difficult to implement across research environments due to the need for specialized laboratory infrastructure and expertise. In contrast, long-term culture is a standardized, accessible method to direct hPSC-CM maturation in a scalable manner that mimics the extended time period in which myocardial maturation occurs *in vivo*^15,17,28–38^. Accordingly, long-term culture is a useful platform to understand the intrinsic molecular determinants of CM maturation and such insights could inform future efforts to accelerate hPSC-CM maturation *in vitro*.

Biologically-inspired approaches to promote hPSC-CM maturation have historically utilized a reductionist framework centered around the quantification of several key maturation phenotypes. More recently, hPSC-CM maturation studies have extended the analytical framework to include -omic profiling most often focusing on the transcriptome, owing to its accessibility and ability to identify molecular pathways and regulatory networks associated with global changes in gene expression during maturation^28,39^. Although studies of hPSC-CM maturation through long-term culture have also independently profiled the proteome^38,40^, metabolome^32,36^, and transcriptome^34,37,41^, few reports have integrated multiple -omic approaches^42,43^ to gain a systems-level view of CM maturation. Despite the clear potential of comprehensive multi-omic approaches for generating mechanistic insights and enabling hypothesis-driven interventions, longitudinal studies tracking hPSC-CM maturation across multiple molecular layers during long-term hPSC-CM culture are severely limited.

To systematically interrogate the mechanisms underlying hPSC-CM maturation during long-term culture, we temporally profiled the metabolome, proteome, and transcriptome from Day 30 to Day 113. Using immunocytochemistry and qPCR-based phenotype assays, we established patterns of structural, sarcomeric, and metabolic maturation. Using both single-omic and integrated multi-omic analytical approaches, we demonstrated that long-term monolayer culture promotes enhanced cardiomyocyte maturation corresponding with molecular changes associated with structural, metabolic, electrophysiological, and cell cycle maturation. Moreover, we also highlight novel signaling pathways and transcription factors accountable for long-term culture duration-dependent hPSC-CM maturation. Furthermore, through several multi-omic analysis approaches, we uncovered unique -omic patterns and pathways underlying duration-dependent hPSC-CM maturation that were obscured from single-omic analyses. By evaluating hPSC-CM multi-omic responses to long-term monolayer culture, this study delineates the intrinsic regulatory programs underlying duration-dependent CM maturation *in vitro* and offers a molecular framework for improving CM maturation for cardiovascular research and therapeutic applications.

## 2. Results

### 2.1 Long-term culture leads to hPSC-CM maturation and design of multi-omic profiling approach

Long-term culture of high purity hPSC-CM cultures has consistently enhanced structural, electrophysiological, and metabolic CM maturation phenotypes^15,17,18,30–33,35–38,43^. For structural maturation, reports have demonstrated that culture as long as 360 days increased sarcomere organization^15,44^ and length^15,30,35,37^ up to ∼1.8-2.0 µm. Similarly, several studies of long-term culture and other hPSC-CM maturation strategies have reported maturation-dependent isoform switches in myosin heavy chain expression at the gene and protein level from MYH6 to MYH7^16,28,38^ as well myosin light chain from MYL7 to MYL2^15,17,32,36,38,43^. For electrophysiological maturation, long-term culture and other hPSC-CM maturation strategies have established improved calcium handling and action potential kinetics^18,30,31^ as well as increases in Cx43 expression^12,30^. Lastly, long-term culture has been shown to impart metabolic changes including elevated oxygen consumption and decreased extracellular acidification^32,36–38^ accompanied by increases in mitochondrial content^32,33^. Although several studies have explored these hPSC-CM maturation phenotypes using single -ome approaches^32,36–38,40,41^, analysis of multi-omes permits understanding coordinate regulation of these phenomena at the gene, protein, and metabolite level.

To investigate hPSC-CM maturation during long-term monolayer culture (**Figure 1A**), we differentiated hPSCs using the GiWi protocol^7,45^ and verified high purity differentiation (mean: 76.2% CMs) using flow cytometry for cTnT+ cells on Day 16 (**Figure 1B, Figure S1A**). Subsequently, we collected parallel hPSC-CM wells on Days 30, 60, and 113 after initiation of differentiation for transcriptomics, proteomics, and metabolomics (**Figure 1A**) as this time window captures hPSC-CMs after lineage specification during a practical and critical period where dynamic changes occur in structural, electrophysiologic, and metabolic maturation phenotypes^18,28,33,35,36^. To validate that parallel wells displayed the same purity as Day 16 wells, we performed CIBERSORTx^46^ analysis on our transcriptome data using the single cell RNA sequencing (scRNAseq) author-defined CM (cTnT+) clusters from Grancharova *et al.* as a reference^41^. CIBERSORTx performs a digital cytometry analysis where a reference scRNAseq dataset is used to predict cell populations from bulk transcriptome data^46^. This analysis suggested no significant difference in the percentage of CMs on Day 16 by flow cytometry compared to long-term culture transcriptome samples (**Figure 1B**). To investigate hPSC-CM maturation in bulk transcriptome samples using CIBERSORTx and the Grancharova scRNAseq reference, we grouped the author-defined CM clusters from Day 12 and Day 24 and labeled them as early CMs and relabeled the Day 90 CM cluster as late CMs. After performing CIBERSORTx digital cytometry on our transcriptome samples, we saw a significant reduction in the percentage of early CMs (**Figure 1C**) and a corresponding increase in the percentage of late CMs (**Figure 1D**) with long-term culture. Together these findings support high CM purity differentiation and provide evidence for transcriptional hPSC-CM maturation in samples collected for multi-omic analysis following long-term culture.

**Figure 1.**
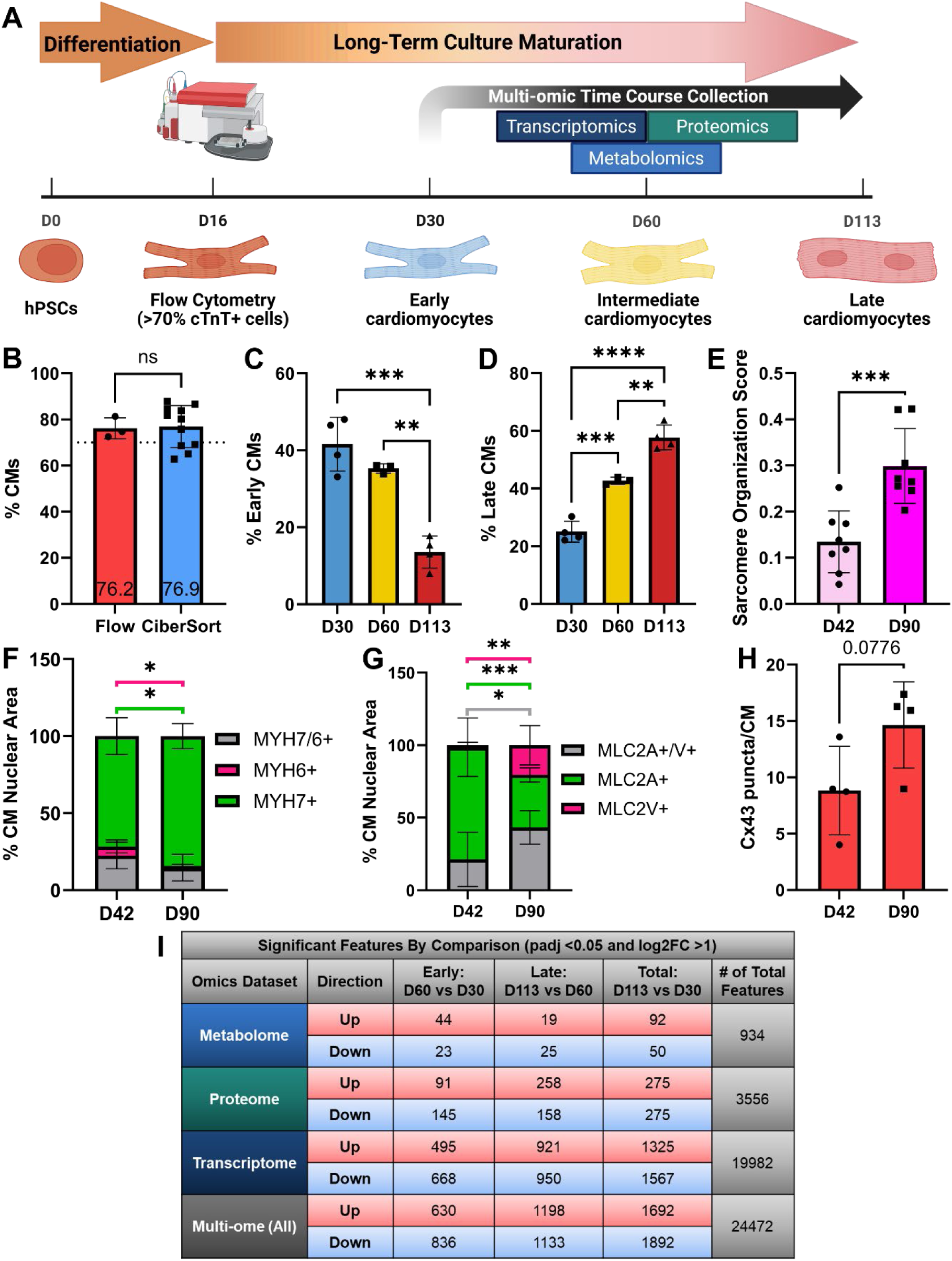
Long-term culture leads to hPSC-CM maturation and design of multi-omic profiling approach. **A)** Overview of multi-omic profiling approach for long-term culture hPSC-CM maturation. **B)** Percentage of cTnT+ cardiomyocytes by flow cytometry on D16 (n=3 replicates) and by CIBERSORTx (CiberSort) digital cytometry transcriptome analysis on D30, D60, and D113 (n=11 replicates, 3-4 per time point). P-value from unpaired t-test. **C)** Percentage of early cTnT+ cardiomyocytes from CiberSort digital cytometry transcriptome analysis (n=3-4 replicates per time point). P-values from one-way ANOVA with Tukey’s post-hoc tests. **D)** Percentage of late cTnT+ cardiomyocytes from CiberSort digital cytometry transcriptome analysis (n=3-4 replicates per time point). P-values from one-way ANOVA with Tukey’s post-hoc tests. **E)** Sarcomere organization score (arbitrary units, range = 0-1) quantified from ACTN2 images using SotaTool at 60x magnification (n=8 images per time point). P-value from unpaired t-test. **F)** Percentage of MYH7+ only, MYH6+ only, and double positive cardiomyocytes based on nuclear area at 20x magnification (n=8 images per time point). P-values from unpaired t-tests (statistical annotation line color corresponds with the legend). **G)** Percentage of MLC2V+ only, MLC2A+ only, and double positive cardiomyocytes based on nuclear area at 20x magnification (n=8 images per time point). P-values from unpaired t-tests (statistical annotation line color corresponds with the legend). **H)** Cx43+ puncta per cardiomyocyte (n=4 images per time point). P-value from unpaired t-test. **I)** Differential feature abundance for each comparison for each - ome for Padj < 0.05 and absolute value of log2FC > 1. P-values for metabolome and proteome from limma. P-values for transcriptome from DESeq2. N=3-4 replicates per time point per -ome.

To verify hPSC-CM maturation with long-term culture, we assayed several phenotypes associated with structural, electrophysiological, and metabolic maturation including sarcomere length^30^, sarcomere organization^44^, myosin heavy and light chain isoform switches^16^, Cx43 gap junction density^12^, and mitochondrial content^33^. To examine structural maturation, we identified sarcomeres by ACTN2 staining and quantified sarcomere length and organization. Using manual and automated (SotaTool^47^) sarcomere length quantification, we noted sarcomere lengths between ∼1.8 and 2.0 µm (depending on the measurement method) on both Day 42 and Day 90 in culture (**Figure S1B-C**). While sarcomere length was unchanged from Day 42 to Day 90, we detected increased sarcomere organization over this time frame using automated quantification^47^ (**Figure 1E**, **Figure S1D**). To further investigate structural phenotypes of hPSC-CM maturation using immunocytochemistry, we examined sarcomere isoform switches by quantifying the percentage of double and single positive CMs for myosin heavy chain markers MYH7 and MYH6 as well as myosin light chain markers MLC2V and MLC2A. For the myosin heavy chain isoform switch from MYH6 to MYH7 (**Figure S1E**), we observed a significant reduction in the single positive MYH6+ CM population and a concomitant significant increase in the single positive MYH7+ CM population from Day 42 to Day 90 (**Figure 1F**). For the myosin light chain isoform switch from MLC2A to MLC2V (**Figure S1F**), we observed a significant increase in the single positive MLC2V+ CM population from Day 42 to Day 90 as expected (**Figure 1G**). Additionally, we observed a significant decrease in the single positive MLC2A+ CM population and an accompanying significant increase in the percentage of double positive MLC2A+/MLC2V+ CMs during long-term culture (**Figure 1G**).

To examine changes in gap junction density during long-term culture, we stained CMs with the MF20 antibody and immunolabeled with an anti-Cx43 antibody to identify gap junctions (**Figure S1G**), then quantified the number of Cx43 puncta per CM. We observed a modest increase in the mean number of Cx43+ puncta per CM following long-term culture (**Figure 1H**). To examine mitochondrial content changes during long-term culture maturation, we measured the ratio of mitochondrial-to-nuclear DNA content using qPCR. Mitochondrial content significantly increased during early maturation from D30 to D60 (**Figure S1H-I**). To summarize, across immunocytochemistry and qPCR measurements we observed multiple lines of evidence for hPSC-CM maturation during long-term monolayer culture, consistent with the literature and our prior work.

To understand the magnitude of how molecular, multi-omic features change during early (D60 vs. D030), late (D113 vs. D60), and total (D113 vs. D30) maturation time course comparisons, we identified significant feature abundance changes using limma^48^ for metabolomics and proteomics and DESeq2^49^ for transcriptomics (**Figure 1I**). Across all -omes, we quantified 24,472 features, including 934 metabolites, 3,556 proteins, and 19,982 transcripts. Comparing D60 vs. D30, 67 metabolites, 236 proteins, and 1,136 transcripts significantly changed. Comparing D113 vs. D60, 44 metabolites, 416 proteins, and 1,871 transcripts significantly changed. Lastly, across the total time course (D113 vs. D30), 142 metabolites, 550 proteins, and 2,892 transcripts significantly changed. Taken together with identified phenotypes of hPSC-CM maturation, these multi-omic trends demonstrate molecular changes during maturation through Day 113 in culture.

### 2.2 Intracellular metabolomic analysis identifies membrane remodeling and fatty acid oxidation as key indicators of hPSC-CM maturation during long-term culture

Untargeted metabolomics identified substantial metabolic rewiring during long-term culture of hPSC-CMs. Principal component analysis (PCA) demonstrated a clear separation between Day 113, Day 60, and Day 30 samples (**Figure 2A**). Hierarchical clustering of sample distances similarly grouped samples by culture duration, with Day 30 samples clustering distinctly from Day 60 and Day 113 samples, demonstrating an early shift in the metabolome (**Figure 2B**). Differential feature abundance analysis identified that 67 of 934 metabolites significantly changed from Day 30 to Day 60 (**Figure S2A**). Consistent with hierarchical clustering, there were fewer significant changes between Day 60 and Day 113 samples, with 44 of 934 metabolites significantly changed (**Figure S2B**). However, metabolic changes during long-term culture clearly increased over time as 142 of 934 metabolites significantly changed from Day 30 to Day 113 (**Figure S2C**). To investigate metabolic pathway-level changes, we were able to map 126 of the 934 metabolites to MetaboAnalyst^50,51^ using HMDB IDs^52^ and identified 20, 7, and 43 significantly changed metabolites during early, late, and total maturation comparisons (D60 vs. D30, D113 vs. D60, and D113 vs. D30, respectively). Due to fewer significantly changed metabolites mapped during early and late maturation comparisons, only 3 pathways were significantly enriched from Day 30 to Day 60, including Phospholipid biosynthesis (**Figure S2D**, SMPDB), Mitochondrial Beta-Oxidation of Long Chain Fatty Acids (**Figure S2E**, SMPDB), and Pentose Phosphate Pathway (**Figure S2F**, KEGG). Phospholipid biosynthesis was driven by a significant increase in phosphatidylglycerol (PG[16:0/16:0]), a direct precursor for cardiolipin, which is necessary for mitochondrial inner membrane oxidative function in highly energy-dependent tissues such as the heart^53^. Moreover, Phospholipid biosynthesis was also driven by a significant increase in phosphatidylinositol (PI[16:0/16:0]), a precursor of phophoinositides, involved in calcium signaling pathways in cardiomyocytes among other roles^54,55^. Upregulation of Mitochondrial Beta-Oxidation of Long Chain Fatty Acids was driven by a significant increase in L-carnitine, which is involved in mitochondrial transport of long-chain fatty acids for ATP production^36,56^. Lastly, increased sedoheptulose drove a significant increase in the Pentose phosphate pathway, likely reflective of glucose shunting towards NADPH production^57^.

**Figure 2.**
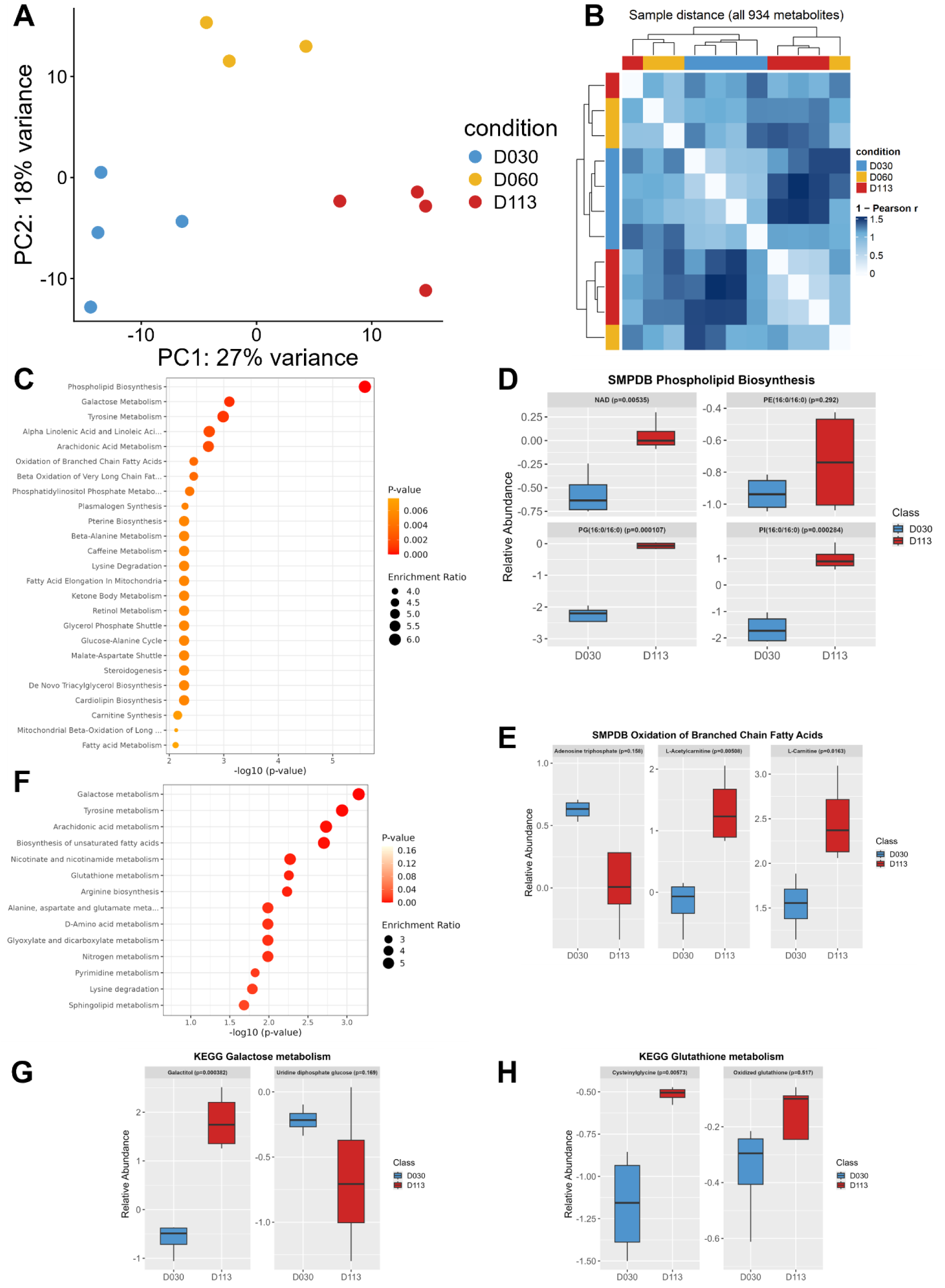
Intracellular metabolomic analysis identifies membrane remodeling and fatty acid oxidation as key indicators of hPSC-CM maturation. **A)** PCA of intracellular metabolite abundances on Day 30, 60, and 113. N=3-4 replicates per time point. Data were log_2_-transformed, autoscaled, and the 500 most variable metabolites across samples were analyzed. **B)** Heatmap of sample distances based on all intracellular metabolites (N=934) calculated using Pearson correlation with lighter colors indicating higher pairwise similarity between samples. **C)** Quantitative enrichment analysis between D113 and D30 displaying the top 25 enriched SMPDB pathways. All pathways shown have FDR-adjusted p < 0.05 and are ordered by raw p-value. **D)** Relative abundance for metabolites identified in the SMPDB Phospholipid Biosynthesis pathway. **E)** Relative abundance of metabolites in the SMPDB Oxidation of Branched Chain Fatty Acids pathway. **F)** Quantitative enrichment analysis between D113 and D30 displaying the 14 enriched KEGG pathways. All pathways shown have FDR-adjusted p < 0.05 and are ordered by raw p-value. **G)** Relative abundance of metabolites in the KEGG Galactose metabolism pathway. **H)** Relative abundance for metabolites in the KEGG Glutathione metabolism pathway. All box plots display log_2_-transformed, normalized metabolite levels for each metabolite at D30 and D113 (N=4 replicates per time point) and p-values are from unpaired t-tests.

Across the total maturation time course (D30 to D113), metabolic pathway remodeling was much more prominent. D30 to D60 increases in Phospholipid biosynthesis (**Figure 2C-D**), Mitochondrial Beta-Oxidation of Long Chain Fatty Acids (**Figure 2C**), and several other terms related to fatty acid oxidation, including Oxidation of Branched Chain Fatty Acids (**Figure 2C, E**), were preserved. In addition to these terms, we also noted significant increases in Galactose Metabolism and Glutathione Metabolism (**Figure 2F**), which were driven by increases in galactitol (**Figure 2G**) and cysteinylglycine (**Figure 2H**) abundances, respectively, and reflect maturation-dependent shifts in energy utilization^58^ and handling of reactive oxygen species^59^. Together, these metabolome findings are consistent with membrane remodeling and increased fatty acid oxidation during long-term hPSC-CM culture.

### 2.3 Proteomic analysis identifies calcium handling and cell cycle exit as key metrics of hPSC-CM maturation

Mass spectrometry-based quantitative proteomics demonstrated extensive changes during long-term culture of hPSC-CMs. PCA established distinct clusters for Day 113, Day 60, and Day 30 samples (**Figure 3A**). Whereas the most noticeable separation in metabolomic profiles occurred between Day 30 and Day 60 samples, hierarchical clustering of proteomic sample distances demonstrated that the most pronounced separation occurred between Day 60 and Day 113 samples, suggesting a later proteomic maturation trajectory (**Figure S3A**). Differential protein abundance identified that 236 of 3556 proteins significantly changed from Day 30 to Day 60 (**Figure S3B**). Consistent with hierarchical clustering, there were more significant changes between Day 60 and Day 113 samples with 416 of 3556 proteins significantly changed (**Figure S3C**). Overall, 550 of 3556 proteins significantly changed across the total time course from Day 30 to Day 113 (**Figure S3D**).

**Figure 3.**
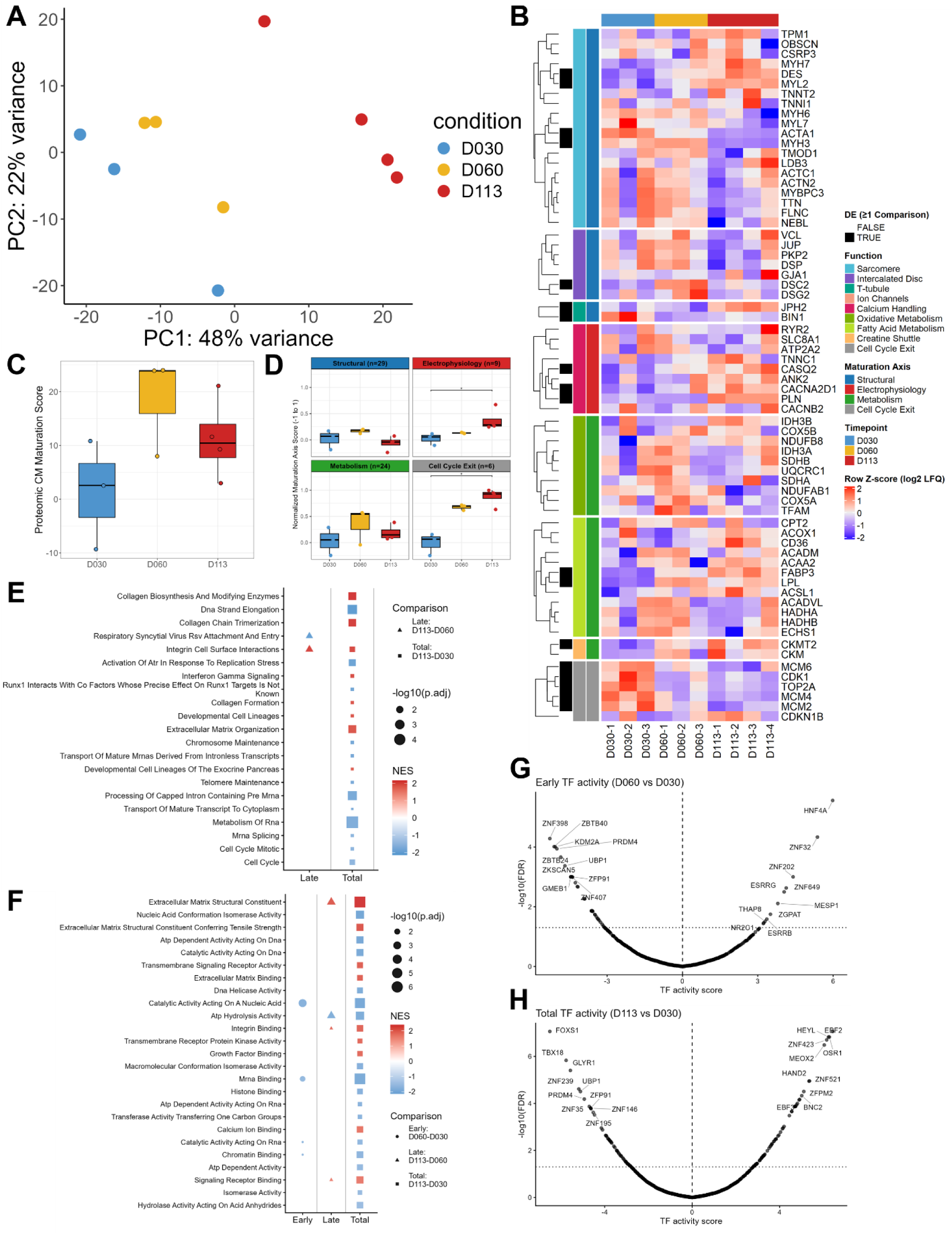
Proteomic analysis identifies calcium handling and cell cycle exit as key metrics of hPSC-CM maturation. **A)** PCA of protein abundances on Day 30, 60, and 113. N=3-4 replicates per time point. Data were log_2_-transformed, autoscaled, and the 500 most variable metabolites across samples were analyzed. **B)** Heatmap of 68 maturation-associated proteins (Row Z-score log2 LFQ). Black squares denote significance in 1 comparison or more. **C)** Proteomic CM maturation score across time in culture. Scores were calculated from signed, capped sample deltas in log2 LFQ intensities summed across all 68 proteins relative to the D30 mean. P-values from Kruskal Wallis with a Dunn’s post hoc test. **D)** Normalized axis-level maturation scores across time in culture. Scores were calculated from signed, capped sample deltas in log2 LFQ intensities and each axis score was normalized by the number of proteins. P-values from Kruskal Wallis with a Dunn’s post hoc test. **E)** Reactome GSEA of proteomic changes. Dot size indicates -log10(padj) and color indicates normalized enrichment score. **F)** GO Molecular Function GSEA of proteomic changes. Dot size indicates -log10(padj) and color indicates normalized enrichment score. **G)** DoRothEA TF activity volcano plot for proteomic early maturation comparison Day 60 vs. Day 30. **H)** DoRothEA TF activity volcano plot for proteomic total maturation comparison Day 113 vs. Day 30. TF activity was inferred using univariate linear modeling (ULM) with Benjamini-Hochberg FDR-adjusted p-values and included all regulons (levels A-E).

To understand the degree of proteomic changes related to maturation during hPSC-CM long-term culture, we curated a list of 113 proteins to describe shifts in key maturation axes, including structural, electrophysiological, metabolism, and cell cycle exit (**Table S1**). We separated these proteins into mature cardiomyocyte functions spanning the sarcomere, intercalated disc, t-tubule, ion channels, calcium handling, oxidative metabolism, fatty acid metabolism, creatine shuttle, and cell cycle exit. From the curated 113 protein list, 68 were detected in at least one sample at each time point, of which 18 were significantly altered during long-term culture in at least one comparison (**Figure 3B**). To summarize the degree of proteomic maturation, we developed a sample-specific proteomic CM maturation score using this targeted protein list. The score was calculated by taking the difference between the log_2_-transformed label-free quantification (LFQ) intensity from the sample of interest and the mean of the Day 30 reference samples for each protein. Subsequently, we capped the delta contribution of each protein at 1 or - 1 and multiplied each protein by the expected direction of change during CM maturation so that proteins expected to increase or decrease could contribute a maximum value of 1 for changes consistent with maturation. Maturation axis and function-level scores were further normalized by the number of proteins in each set to account for differences in set size across maturation modules of interest. Overall, late-stage increases in the proteomic CM maturation score (**Figure 3C**) were primarily driven by changes in proteins related to electrophysiology and cell cycle exit from Day 30 to Day 113 (**Figure 3D**). Because no ion channels were detected, electrophysiological maturation was driven by significant increases in calcium handling proteins including CASQ2, CACNA2D1, and PLN (**Figure S3E**). While structural (33 proteins) and sarcomere maturation scores (20 proteins) did not increase over this time frame, we observed significant increases in sarcomere isoform protein ratios with the MYH7/MYH6 (**Figure S3F**) and MYL2/MYL7 ratios (**Figure S3G**) rising significantly across the long-term culture time course. Other notable structural protein changes included a significant increase in DES and a decrease in ACTA2. Additionally, there were significant increases in fatty acid metabolism proteins LPL and FABP3 as well as creatine shuttle protein CKMT2 (**Figure 3B**).

To profile proteomic changes in an unbiased way, we performed GSEA analyses^60^ for multiple pathway databases as well as transcription factor (TF) regulon analyses^61^ across early (D60 vs. D30), late (D113 vs. D60), and total (D113 vs. D30) time periods. Using Reactome GSEA analysis, we observed increased Extracellular Matrix Organization and decreased Cell Cycle over the total maturation time course (**Figure 3E**). Similarly, using GO Molecular Function GSEA analysis, we observed significant increases in several ECM-related terms during late and total maturation comparisons (**Figure 3F**). Additionally, consistent with targeted proteome analyses regarding calcium handling, we observed a significant increase in Calcium Ion Binding over the entire maturation time course (**Figure 3F**). Lastly, for Hallmark GSEA analysis, we observed a significant increase in Myogenesis during the total maturation time course (**Figure S3H**). For TF regulon analyses, we observed significant increases in ESSRG, ESSRB, and MESP1 TF activity from Day 30 to Day 60 (**Figure 3G**). Moreover, during late (**Figure S3I**) and total (**Figure 3H**) maturation comparisons, we observed consistent, significant increases in HEYL, OSR1, MEOX2, EBF2, and ZNF423. Moreover, from Day 30 to Day 113 we also noted a significant increase in HAND2 activity (**Figure SH**). Together, these inferred TF activities suggest a role for estrogen signaling (ESRRG and ESRRB)^62,63^, Notch signaling (HEYL)^64^, and mesodermal developmental TFs (MESP1, MEOX2, OSR1)^65,66^ during hPSC-CM maturation. Further, we identified potential novel hPSC-CM maturation TFs, including EBF2 and ZNF423, which are known to be important for mitochondrial function^67,68^.

### 2.4 Transcriptomic analysis identifies improvements in electrophysiology, metabolism, and cell cycle exit during long-term culture hPSC-CM maturation

Transcriptomics displayed more consistent temporal changes than metabolomics and proteomics throughout long-term hPSC-CM culture. Accordingly, Day 113, Day 60, and Day 30 samples displayed clear separation across both PCA (**Figure 4A**) and hierarchical clustering of sample distances (**Figure S4A**). Notably, 1163 of 19982 transcripts significantly changed from Day 30 to Day 60 (**Figure S4B**), 1871 of 19982 transcripts significantly changed from D60 to Day 113 (**Figure S4C**), and 2892 of 19982 significantly changed from Day 30 to Day 113 (**Figure S4D**).

**Figure 4.**
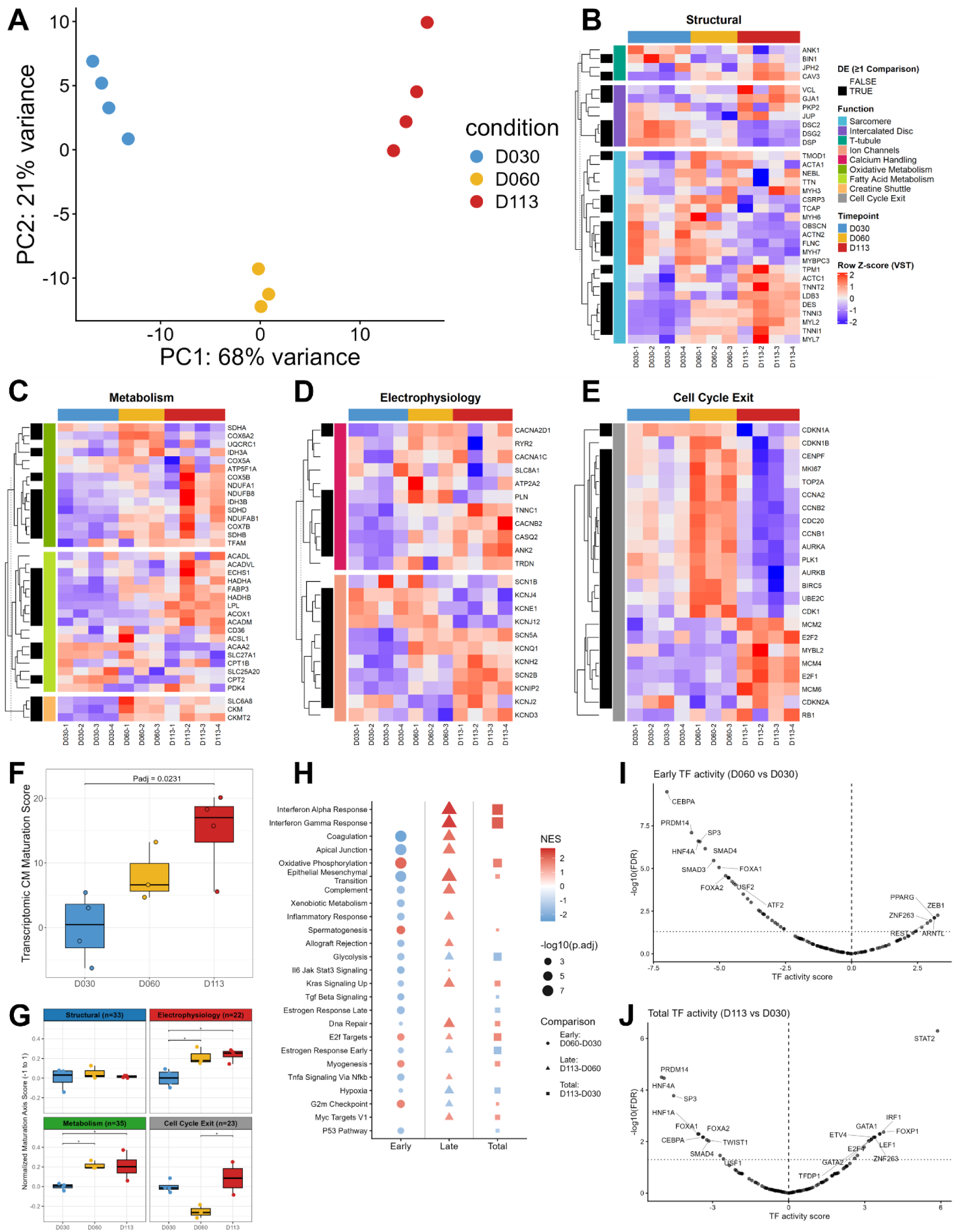
Transcriptomic analysis identifies improvements in electrophysiology, metabolism, and cell cycle exit during long-term culture hPSC-CM maturation. **A)** PCA of transcript abundances on Day 30, 60, and 113. N=3-4 replicates per time point. Data were VST-transformed, autoscaled, and the 500 most variable metabolites across samples were analyzed. **B-E)** Heatmaps of 113 maturation-associated transcripts (Row Z-score VST). Black squares denote significance in 1 comparison or more. **B)** Heatmap for Structural maturation axis. **C)** Heatmap for Metabolism maturation axis. **D)** Heatmap for Electrophysiology maturation axis. **E)** Heatmap for Cell Cycle Exit maturation axis. **F)** Transcriptomic CM maturation score across time in culture. Scores were calculated from signed, capped sample deltas in VST-transformed counts summed across all 113 transcripts relative to the D30 mean. P-values from Kruskal Wallis with a Dunn’s post hoc test. **G)** Normalized axis-level maturation scores across time in culture. Scores were calculated from signed, capped sample deltas in VST-transformed counts and each axis score was normalized by the number of transcripts. P-values from Kruskal Wallis with a Dunn’s post hoc test. **H)** Reactome GSEA of transcriptomic changes. Dot size indicates - log10(padj) and color indicates normalized enrichment score. **I)** DoRothEA TF activity volcano plot for transcriptomic early maturation comparison Day 60 vs. Day 30. **J)** DoRothEA TF activity volcano plot for transcriptomic total maturation comparison Day 113 vs. Day 30. TF activity was inferred using univariate linear modeling (ULM) with Benjamini-Hochberg FDR-adjusted p-values and restricted to high-confidence regulons (levels A and B).

To profile long-term culture hPSC-CM transcriptomic changes across multiple maturation axes, we used the same curated list evaluated using proteomics to assay the expression of 113 genes involved structural (**Figure 4B**, 33 genes), metabolism (**Figure 4C**, 35 genes), electrophysiology (**Figure 4D**, 22 genes), and cell cycle exit (**Figure 4E**, 23 genes) maturation axes. All 113 genes were detected in transcriptomics with widespread changes across all maturation axes; 22/33 structural genes, 23/35 metabolism genes, 15/22 electrophysiology genes, and 21/23 cell cycle exit genes significantly changed in at least one comparison. The transcriptomic CM maturation score, which was calculated using an analogous approach to proteomics starting with variance stabilizing transformation (VST)-transformed counts^49^, significantly increased between Day 30 and Day 113 (**Figure 4F**), driven by early increases in expression of genes involved in electrophysiology and metabolism maturation and a later increase in cell cycle exit genes (**Figure 4G**). While the structural axis-level maturation score did not significantly increase, there was a late-stage increase in the t-tubule function-level maturation score without maturation-consistent changes across sarcomere and intercalated disc function-level maturation scores (**Figure S4E**). Moreover, we observed a strong temporal increase in the *MYL2/MYL7* gene expression ratio (**Figure S4F-G**), consistent with protein-level findings of sarcomere maturation. The improvement in electrophysiological maturation was driven by late-stage increases in ion channels and early-stage increases in calcium handling (**Figure S4E**). Similarly, the improvement in the metabolism maturation axis was driven by late-stage increases in fatty acid metabolism-related genes and early-stage increases in creatine shuttle and oxidative metabolism-related genes (**Figure S4E**). Together, these findings suggest time-dependent changes in maturation where significant portions of metabolic, electrophysiologic, and specific portions of structural maturation occurred concurrently across this time window and preceded late-stage cell cycle exit.

To further assess long-term culture-dependent hPSC-CM transcriptomic changes using discovery-based approaches, we quantified pathway and TF activity using GSEA^60^ and TF regulon^61^ analyses. From GSEA, we noted pathway changes consistent with our targeted analysis, including increased Oxidative Phosphorylation, Myogenesis, and G2M Checkpoint as well as decreased Glycolysis (**Figure 4H**). Similarly, we observed increases in the Respiratory Chain Complex (**Figure S4H**), Muscle Contraction (**Figure S4I**), and Respiratory Electron Transport (**Figure S4J**). Beyond these findings, we observed decreases in TGFβ signaling (**Figure 4H**), ERK1 and ERK2 Cascade (MAPK signaling)^69^, and the development of other organ systems (**Figure S4I)**. Moreover, there were significant increases in Proteasome Complex, Spliceosomal Complex (**Figure S4H**), and mRNA splicing (**Figure S4J**), which suggests increased regulation of protein turnover^37^ and transcript processing^70,71^. To increase the specificity of TF activity inference from the large transcriptome feature space, we restricted our analysis to high-confidence DoRothEA regulons (levels A-B) to yield a more interpretable set of regulators. Using this approach, we noted significant a decrease in SMAD3 TF activity during long-term hPSC-CM culture, which was consistent with pathway analyses demonstrating decreased TGFβ signaling^72^ (**Figure 4I**). Additionally, we observed strong decreases in activity of TFs known to be involved in the development of other organ systems or the maintenance of pluripotency, including decreases in FOXA1/2^73^, HNF1A/4A^74^, CEBPA^75^, and PRDM14^76^ activity (**Figure 4I-J, Figure S4K**). During early hPSC-CM maturation (D30 to D60), we observed significant increases in PPARG and ARNTL activity^77^, consistent with early improvements in metabolism. From Day 60 to Day 113, we inferred significant increases in AP1 transcription factors FOSL1 and JUN, which have been reported to regulate murine postnatal heart maturation^78–80^. During the total hPSC-CM maturation time course (D30 to D113), there were significant increases in E2F4 and TFDP1 activity, which are regulators of cell cycle exit^81^ (**Figure 4J**). On the whole, these discovery-based pathway and TF activity analyses verify significant findings from targeted transcriptome analysis and further emphasize potential novel drivers of hPSC-CM maturation during long-term culture.

### 2.5 Integrated multi-omic and machine learning analyses identifies novel time-dependent maturation programs

To discover cross-layer biological patterns and interactions during long-term hPSC-CM culture, we employed several multi-omic integration strategies. Foremost, we utilized Multi-omic Factor Analysis (MOFA)^82,83^, an unsupervised machine learning approach, to discover multi-omic patterns in the data. We used all 934 metabolites, all 3556 proteins, and the top 5000 most variable transcripts to force a similar magnitude of features across -omes. Factors 1 and 2 explained the largest degree of shared multi-omic variance, and were able to confidently distinguish D30, D60, and D113 hPSC-CM samples using multi-omic factor plot analysis for dimensionality reduction (**Figure 5A**). Moreover, Factor 1 captured the dominant source of expected variation in the data, with sample factor values significantly increasing with time in culture (**Figure 5B**). Across all 6 factors, 85.3% of transcriptomic variance, 73.9% of proteomic variance, and 31.0% of metabolomic variance was explained (**Figure S5A**). Factor 1 exhibited the strongest signal from the transcriptome and captured most of the variance across all 6 factors with 58.0% of transcriptomic variance, 32.9% of proteomic variance, and 19.4% of metabolomic variance represented (**Figure 5C**). Factor 2 variance was more evenly distributed among -omes with 17.9% of transcriptomic variance, 11.7% of proteomic variance, and 9.2% of metabolomic variance represented (**Figure 5C**). Notably, Factor 3 explained less than 2% of the transcriptomic and metabolomic variance but captured 25.0% of the proteomic variance while Factors 4 through 6 explained less than 10% of any -omic variance summed across all three factors.

**Figure 5.**
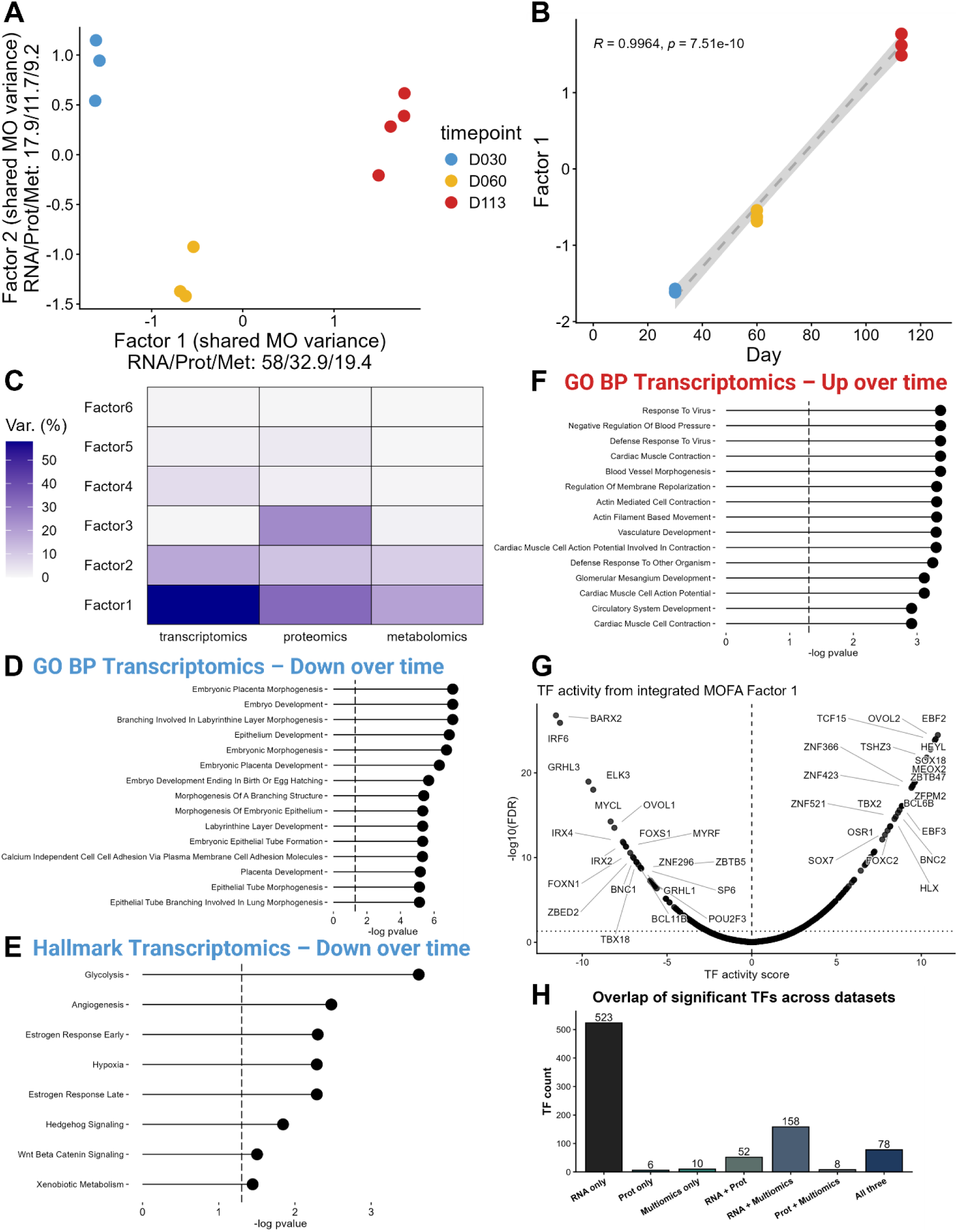
Integrated multi-omic and machine learning analyses identifies novel time-dependent maturation programs. **A)** Multi-omic factor plot of sample factor values for MOFA factors 1 and 2 for D30, D60, and D113 samples (N=3-4 replicates per time point). MOFA was performed with all 934 metabolites, all 3556 proteins, and the top 5000 most variable transcripts. **B)** Factor 1 sample factor values over time in culture. P-value and R-value from linear regression. **C)** Percent variance explained by each factor across -omes. **D)** GO BP GSEA for negative transcriptome factor values. **E)** Hallmark GSEA for negative transcriptome factor values. **F)** GO BP GSEA for positive transcriptome factor values. **G)** DoRothEA TF activity volcano plot from integrated MOFA Factor 1 weights (sum of transcriptomic and proteomic weights). **H)** Summary of the number of significant TFs unique and shared across -omic layers. RNA = transcriptome. Prot = Proteome. TF activity was inferred using univariate linear modeling (ULM) with Benjamini-Hochberg FDR-adjusted p-values and included all regulons (levels A-E) for all -omes.

Because sample Factor 1 values progressively increased with time in culture, features (metabolites, proteins, or transcripts) with positive weights were interpreted as progressively increasing in abundance with time in culture and features with negative weights were interpreted as progressively decreasing in abundance with time in culture. Therefore, to investigate time-dependent hPSC-CM maturation, we explored biological pathways that changed consistently with time in culture by performing enrichment analyses separately for features with negative and positive Factor 1 weights. For transcripts with negative Factor 1 weights, which decreased significantly across the time course, we discovered GO BP GSEA pathways that were primarily related to embryonic, placental, or epithelial development (**Figure 5D**). Similarly, for transcripts decreasing with time, Hallmark GSEA pathway analysis indicated significant decreases in Glycolysis, Hedgehog Signaling, and Wnt Beta Catenin Signaling (**Figure 5E**). For the proteins significantly decreasing with time, Hallmark GSEA pathway analysis also demonstrated E2F targets and MYC targets as significant (**Figure S5B**), consistent with decreased cell cycle progression^84^. To understand molecular programs that increased with hPSC-CM maturation, we investigated features with positive Factor 1 weights. Using GO BP GSEA pathway analysis of gene expression, we found pathways increasing with time in culture such as Cardiac Muscle Contraction, Regulation of Membrane Repolarization, Actin Mediated Cell Contraction, and Cardiac Muscle Cell Action Potential (**Figure 5F**). Similarly, for proteins that increased with time in culture, GO MF GSEA pathway analysis demonstrated clear increases in Calcium Ion Binding and several ECM-related terms (**Figure S5C**). In total, time-dependent pathway analyses suggest key changes consistent with cardiomyocyte maturation including improvements in terms-related to metabolism^19^, action potential^18^, contraction, and cell cycle exit^85^. Moreover, downregulated transcripts with time suggest Wnt^86,87^ and Hedgehog^88,89^ signaling inhibition as potential intrinsic time-dependent mechanisms of hPSC-CM maturation with long-term culture.

Beyond signaling pathway changes, we investigated multi-omic temporal TF activity changes using Factor 1 weights from MOFA analysis from both the proteome and transcriptome. To integrate the proteome and transcriptome, Factor 1 weights were summed across both -omes. By doing this, gene-protein pairs with concordant temporal changes became more heavily weighted while gene-protein pairs with discordant changes became less heavily weighted. Moreover, genes or proteins that were only present in a single -ome were penalized by not having potential additive effects from both -omes. Using this approach, we confirmed several TFs from individual -omic analyses such as novel cardiomyocyte candidate TFs EBF2 and ZNF423 as well as HEYL (Notch signaling), MEOX2, and OSR1 (mesodermal development TFs) (**Figure 5G**). To prioritize TFs across all DoRothEA regulons (levels A-E), we overlapped the significant regulators from the integrated multi-omic analysis of Factor 1 with the significant regulators from any temporal comparison (Day 30 to Day 60, Day 60 to Day 113, Day 30 to Day 113) in both the proteome and transcriptome analyses (**Figure 5H**). Interestingly, we identified 78 TFs that were significant across all 3 analyses (**Figure S5D**) and 10 TFs that were only significant in the multi-omic analysis (**Figure S5E**). In contrast, 747 TFs were significant across all other analysis groups, highlighting the inherent difficulty in prioritizing TF candidates using single -ome or dual analysis overlap approaches.

Using this multi-omic overlap approach to prioritize TFs, we identified several TFs shared across all 3 analyses (**Figure 5G**, **Figure S5D**) that were enriched with time in culture including SOX7 and SOX18^90,91^, which are collectively known to be upregulated by GATA4 during early cardiogenesis and inhibit Wnt signaling in model organisms. While their role in cardiomyocyte fate commitment is well understood in model organisms, evidence for regulating human CM differentiation or maturation is limited. We also observed increased TBX2 activity (**Figure 5G**, **Figure S5D**) with long-term hPSC-CM culture, which is an underexplored paralog of TBX5^92,93^ and TBX20^94,95^ that are known to regulate CM development and postnatal functions. Recently, a study reported that TBX2 is also upregulated by GATA4 specifically in hPSC-CMs during early differentiation^96^. Moreover, ZFPM2 (better known as FOG2) activity, was significantly upregulated with long-term culture (**Figure 5G**, **Figure S5D**). Interesting, ZFPM2 is also known to interact with GATA family transcription factors including GATA4 and displays clear roles in cardiomyocyte development^97^ and postnatal ventricular function^98^ in mice. Beyond these GATA4-interacting transcription factors, we observed a significant increase in TCF15 activity (**Figure 5G**, **Figure S5D**) and a significant decrease in MYCL activity (**Figure 5G**), which are consistent with increased fatty acid uptake^99^ and decreased cell cycle progression^100^ respectively. Additionally, we identified 10 TFs that were uniquely significant in the multi-omic TF analysis including POU3F2, STAT1, and BACH2, which increased with long-term culture (**Figure S5E**) and have relatively unknown roles in CM maturation. Together, our integrated TF activity analysis prioritizes several GATA4-dependent TFs with likely roles in CM maturation during long-term culture, provides support for previously identified maturation phenotypes, and uniquely identifies a set of 10 TFs associated with long-term culture maturation.

To better understand specific -omic features that changed with time, we examined metabolites, proteins, and transcripts with negative and positive Factor 1 weights and assessed gene-protein pairs with conserved changes across time. Out of the top 10 metabolites with highest absolute Factor 1 weights, two phosphatidylinositol species and galactitol significantly increased with time (**Figure S6A**). For proteomics, we observed significant increases in ECM-related proteins EMILIN1 and COL6A3^101^ as well as LGALS1^102^, which has been shown to colocalize with I bands in cardiac sarcomeres. Additionally, we noted increased SFRP5^103^, which is a known Wnt signaling inhibitor (**Figure S6B**). For transcriptomics, we observed significant increases in FGF12^104^ and LPL^105^, which demonstrate important signaling and metabolic features of hPSC-CM maturation (**Figure S6C**). During early maturation from D30 to D60, we observed significant gene-protein pair increases in several heat shock-related proteins which are known to be important for cardiac muscle function,^29^ including HSPB6^106^, CRYAA, CRYBB1, and CRYBB2 (**Figure S6D**). In late maturation from D60 to Day 113, we observed significant gene-protein pair increases in several ECM proteins as well as a significant decrease in LIN28A^107^, a known regulator of pluripotency maintenance (**Figure S6E**). Across the maturation time course from D30 to Day 113, we also observed significant increases in CPNE5^108^, a calcium-dependent membrane protein, and LPL^105^, which breaks down triglycerides (**Figure S6F**). Notably, we saw the greatest conservation of changes across the transcriptome and proteome for gene-protein pairs concordantly upregulated in late (D113 vs. D60, 40 features) and total (D113 vs. D30, 60 features) maturation comparisons. Using GO BP ORA pathway analysis, we noted significant increases in Cell Adhesion, Circulatory System Development, Muscle Contraction, and Regulation of Monoatomic Ion Transport across these comparisons (**Figure S7A-B**).

To extend the integrated pathway analysis beyond the transcriptome and proteome, we performed MultiGSEA^109^, which allows rank-based pathway enrichment across 18009 transcripts, 3374 proteins, and 150 HMDB-annotated metabolites. For WikiPathways MultiGSEA, Fatty acid beta-oxidation was significantly enriched over the total time course (D113 vs. D30) and was driven primarily from metabolites (**Figure S7C**). In late maturation (D113 vs. D60), Panther MultiGSEA identified a significant increase in integrin signaling^110^ (proteome-driven), which has been shown to be involved in hPSC-CM maturation, as well as Slit/Robo signaling^111^ (**Figure S7D,** transcriptome-driven). KEGG MultiGSEA identified a significant increase in the regulation of actin cytoskeleton (**Figure S7E,** transcriptome-driven). Lastly, Reactome MultiGSEA identified significant increases in several mRNA splicing^70,71,112^ terms during late and total maturation comparisons (proteome- and transcriptome-driven). Additionally, we observed a significant increase in aerobic respiration and respiratory electron transport during early maturation, which was driven by transcriptome signals (**Figure S7F**). Together, our MultiGSEA results provide an integrated pathway view and demonstrate several metabolism, structural, and other maturation-associated signals across the multi-ome.

## 3. Discussion

Defining the coordinated molecular regulation of CM maturation remains critical for improving the utility of hPSC-CMs across cardiovascular research and translational applications^2,12^. Here, we leveraged long-term hPSC-CM culture to map the integrated multi-omic trajectory of hPSC-CM maturation during long-term culture *in vitro*. We observed several key maturation hallmarks, including myosin heavy and light chain isoform switches, enhanced sarcomere organization, increased gap junction density, and early mitochondrial biogenesis, over the 84 day time course of this study. Across the multi-omic landscape, asynchronous temporal trends in maturation were apparent, characterized by early functional shifts in the metabolome (Day 30 to Day 60), prominent later-stage shifts in the proteome (Day 60 to Day 113), and steady transcriptome remodeling throughout long-term culture. Metabolomics revealed increases in metabolites related to membrane composition and mitochondrial oxidative function, which corresponded with increases in mitochondrial content. Proteomics identified consistent increases in mature sarcomere isoform ratios and later-stage increases in calcium handling and cell cycle exit, suggesting a metabolic priming effect for specific CM maturation phenotypes. Proteomic pathway-based analyses suggested a consistent role for ECM and integrin-related signaling in regulating CM maturation, and TF activity analyses suggested that estrogen-related receptors (ESRRG and ESRRB), Notch (HEYL) signaling, and several canonical mesoderm developmental TFs may contribute to CM maturation regulation. Transcriptome analyses demonstrated strong electrophysiological, metabolic, and cell cycle exit maturation signatures, while discovery-based analyses suggested signaling pathway (MAPK and TGFβ inhibition) and transcription factor (PPARG and ARNTL) mechanisms for hPSC-CM maturation. Importantly, integrated multi-omic analyses supported these findings and uncovered further compelling CM maturation mechanisms involving Wnt, Sonic Hedgehog, and Slit/Robo signaling for future investigation, as well as time-dependent candidate maturation markers across the metabolome, proteome, and transcriptome.

Despite advances in directing hPSCs to CMs *in vitro*, current directed differentiation protocols largely generate phenotypically immature CMs in comparison to adult CMs^113^. Due the limited availability of human tissue during critical late gestation and early postnatal CM maturation windows^113,114^, our current mechanistic understanding of maturation relies heavily on mouse models, which demonstrate clear differences in maturation timing^115,116^ as well as a fundamental reversal of the expression pattern of canonical MYH7/MYH6 sarcomere isoforms^113,116^. As such, hPSC-CMs represent a powerful model system to elucidate the control of human cardiogenesis and investigate mechanistic hypotheses for promoting human CM maturation. In isolated proteome and transcriptome analyses we discovered several interesting signaling pathway candidates for potentially stimulating hPSC-CM maturation, including increased ECM binding and integrin-related signaling^110,117,118^, TGFβ inhibition^72,119^, and MAPK inhibition^69^. In support of these findings, recent studies demonstrate a role for collagen-based integrin stimulation^110^ and adult human heart decellularized ECM^117^ containing LGALS1 and COL6 in accelerating maturation in hPSC-CM models. Using a small molecule approach to achieve dual MAPK and PI3K/AKT inhibition^69^, Garay *et al*. demonstrated improvements in structural, metabolic, electrophysiology, and cell cycle exit measures of hPSC-CM maturation. While direct evidence for TGFβ inhibition in facilitating CM maturation is limited, the authors also noted concomitant decreases in TGFβ signaling as a result of their dual inhibition approach. Moreover, active TGFβ/SMAD3 signaling during zebrafish cardiac regeneration is required for cell cycle reactivation^72^ implying that TGFβ/SMAD3 inhibition may be involved in regulating cell cycle exit, which is a well-recognized hallmark of mature human CMs.

Surprisingly, while integrated multi-omic analyses substantiated increased integrin-related signaling during maturation, they also uniquely revealed putative functions for Wnt inhibition^86,87,120^, Hedgehog inhibition^88,89,121,122^, and Slit/Robo activation^111^ in guiding hPSC-CM maturation. While Wnt signaling inhibition decisively drives mesoderm specification towards cardiac mesoderm and ultimately CMs^45^, its function in explicitly advancing human CM maturation is underexplored. Seminal work by Buikema and colleagues demonstrated that reactivation of canonical Wnt signaling (via GSK3β inhibition) keeps hPSC-CMs in an immature state and enables rapid CM proliferation^86^. Moreover, several studies have linked the specific timing^87^ and mode^120^ of Wnt inhibition during hPSC-CM differentiation to changes in structural, metabolic, electrophysiological and cell cycle maturation phenotypes. Interestingly, this observation suggests that signaling pathways implicated in CM specification may also influence maturation programs, consistent with reports linking inhibition of MAPK and TGFβ signaling to both CM differentiation^123–125^ and maturation^69,72^. In line with this potential dual inhibition, multi-omic analyses also indicated a role for Hedgehog signaling inhibition in hPSC-CM maturation, which has been shown to promote CM cell cycle exit^88,89^ as well as CM differentiation^121,122^ from cardiac progenitors across hPSC-CM and mouse developmental models. Lastly, integrated multi-omic pathway analysis with MultiGSEA^109^ revealed a prospective role for Slit/Robo activation in hPSC-CM maturation during long-term culture. Evidence for Slit/Robo signaling during human CM development and maturation is particularly sparse, with existing literature supporting Slit/Robo activation in controlling cardiac chamber morphogenesis as well as cardiomyocyte polarity and alignment in model organisms, processes that may set the stage for structural maturation^111^. Taken together, our multi-omic results suggest novel signaling pathways involved in hPSC-CM maturation and differentiation, representing an open area for targeted future investigations.

Although signaling pathways provide important upstream regulatory cues, hPSC-CM maturation is ultimately likely governed by downstream transcription factor-directed gene regulatory networks. Using inferred TF activities from the proteome and transcriptome, we detected several established, emerging, and candidate regulators of CM maturation. From late-stage (D60 to D113) transcriptome analyses, we identified increased FOSL1 and JUN activity, which are AP-1 transcription factors that drive murine postnatal CM maturation across functional domains^79,80^ and are known to be deficient in hPSC-CM maturation systems^78^. Additionally, increases in metabolism-related TFs PPARG and ARNTL (BMAL1) correlated with our observation of early metabolic maturation during long-term culture across both transcriptome and metabolome analyses. While PPARA^126,127^ and PPARD^128^ are well-known regulators of hPSC-CM maturation, PPARG^129^, which is often reported to control lipid metabolism, remains relatively unexplored in maturation contexts. Moreover, PPARG interacts with circadian rhythm transcriptional machinery^129^. Accordingly, we observed increases in circadian clock TF ARNTL (BMAL1) activity in transcriptome analysis, which has been implicated in metabolic maturation of hPSC-CMs^130^. For proteome-inferred TF activities, we uncovered hPSC-CM maturation roles for estrogen-(ESRRG and ESRRB) and Notch-related (HEYL) TFs as well as canonical mesodermal developmental TFs (MESP1, MEOX2, and OSR1). Notably, estrogen-related TFs^62,63,131^ play roles in cardiac metabolic maturation while HEYL (Notch signaling) exhibits temporally-dependent maturation effects with detrimental effects postnatally owing to increased cell cycle progression and positive effects in accelerating CM maturation while activated during hPSC-CM differentiation^132^. In further support of the proposed dual function of developmental signaling and transcriptional programs in both CM differentiation and maturation, we observed increased activity of mesoderm-associated TFs MESP1, MEOX2 and OSR1 during long-term hPSC-CM maturation, consistent with reports showing that mesodermal priming by TBXT can influence downstream maturation programs^66^. Lastly, we identified potential novel hPSC-CM maturation functions for EBF2, a mitochondrial biogenesis regulator^67^, and ZNF423^68^, both of which are known to interact with PPARG. On the whole, these findings demonstrate established roles and suggest novel mechanisms of TF control of CM maturation.

After profiling time-dependent patterns across the multi-ome using MOFA, we further integrated transcriptome and proteome MOFA Factor 1 weights to infer temporal multi-omic TF activity. Notably, several TFs from our analysis supported CM maturation functions including fatty acid uptake^99^ (TCF15 increased with time) and cell cycle exit^100^ (MYCL decreased with time). However, our multi-omic TF activity analysis also prioritized a set of GATA4-dependent TF that increased during long-term culture CM maturation including SOX7^90,91^, SOX18^90,91^, TBX2^96^, and ZFPM2 (FOG2)^97,98^. To date, this core set of GATA4-dependent transcription factors is understood to regulate cardiomyocyte differentiation across different developmental models. In mice, ZFPM2 (FOG2) has demonstrated importance in postnatal ventricular function^98^, and SOX7 expression has recently been reported to increase during postnatal CM maturation^80^. While specific knowledge of these TFs during human CM maturation is limited, their importance during CM lineage commitment across models of CM development is consistent with the parallel changes for developmental TFs and signaling pathways observed across our analyses. Collectively, multi-omic TF activity analyses delineate a core network of GATA4-dependent TFs, namely SOX7, SOX18, TBX2, and ZFPM2 (FOG2), that increase during hPSC-CM maturation and are not clearly prioritized by single -ome analyses. Moreover, multi-omic analysis uniquely identified increased STAT1, BACH2, POU3F2 activity during long-term hPSC-CM culture. While STAT1^133^ and BACH2^134^ have been shown to reduce pathological cardiac hypertrophy in mouse models, the potential role of STAT1, BACH2, and POU3F2 in hPSC-CM maturation remains underexplored.

While TFs direct CM maturation through gene regulatory networks, our multi-omic analyses also revealed several downstream molecular markers reflecting long-term culture hPSC-CM maturation. Most notably, we observed temporal increases in LGALS1^102^, a canonical ECM protein known to interact with I bands in cardiac sarcomeres, across all time points for both the proteome and transcriptome. Moreover, we also noted consistent temporal increases in several ECM proteins during maturation including EMILIN1^135^, COL6A3^117^, and COL1A1^110^ across the proteome and transcriptome, which have demonstrated roles in CM structural maturation. Additionally, LPL^136^, which liberates fatty acids for energy production, was consistently upregulated across the proteome and transcriptome during long-term hPSC-CM culture. Consistent with pathway findings of Wnt inhibition during long-term CM culture, we observed dynamic increases in SFRP5^137^, a secreted antagonist of Wnt signaling, across the proteome and transcriptome. Finally, long-term hPSC-CM culture resulted in enhanced expression of several heat shock-related proteins including CRYAB^29^, CRYAA, CRYBB1/2, and HSPB6^106^, suggesting the importance of protein homeostasis in CM maturation. In sum, through multi-ome analyses we unveiled unappreciated markers of hPSC-CM maturation during long-term culture *in vitro*.

In conclusion, our study highlights how an integrated multi-omic assessment of hPSC-CM maturation during long-term culture can deliver rich, novel insights and provide important context for the pace and degree of maturation across key CM functional domains. Moreover, our results reinforce the utility of long-term culture as a robust strategy to mature hPSC-CMs *in vitro.* Interestingly, our analyses connect signaling pathways and TFs known to strengthen CM lineage commitment with hPSC-CM maturation during long-term culture. Our in-depth analytical approach further provides candidate multi-omic markers of CM maturation, which will serve as a useful reference for future studies. Subsequent investigations should evaluate the intrinsic time-dependent signaling- and TF-based mechanisms described here to accelerate *in vitro* CM maturation. Further, future studies should apply multi-omic characterization to better delineate hPSC-CM maturation dynamics in specialized, biologically-inspired maturation systems.

## 4. Materials and Methods

### 4.1 Human Pluripotent Stem Cell Maintenance

WTC11 hiPSCs were cultured in mTeSR1 (STEMCELL Technologies 85850) on 6-well plates (Corning COSTAR 07-200-82) coated with Growth Factor Reduced Matrigel (0.5 mg/plate or 11 μg/cm2, Corning 354230)^7,138^. hPSCs were maintained in a cell culture incubator (Sanyo MCO-18AC; 37°C, 5% CO2, 95% RH) and passaged with Versene (Life Technologies 15040066) every 3-5 days at 60-80% confluency.

### 4.2 hPSC-Cardiomyocyte Differentiation and Long-Term Culture Maturation

As previously described, hPSCs were lifted on Day -2 for 5 min with Accutase (Innovative Cell Technology AT104) and quenched 1:1 v/v in DMEM/F12 (ThermoFisher 11330032)^7,138^. hPSCs were resuspended in mTeSR1 with 5 μM Y-27632 (Tocris 1254) and seeded onto Matrigel-coated 12-well plates in 2 mL of media at optimal seeding densities for WTC11 hiPSCs. To promote uniform cell attachment, plates were set at room temperature for 30 min prior to transfer to a cell culture incubator. On Day −1, an mTesR1 medium change was performed. On Day 0, differentiation induction was performed in RPMI1640 medium (Life Technologies 11875119) with 2% v/v B27 minus insulin (B27-) and CHIR99021 (Selleckchem S1263) at an optimal concentration between 6-12 μM. Media was replaced with B27- and 5 μM IWP2 (Tocris 3533) on Day 2 and with B27-only on Day 4. On Days 6, 8, 10, and 13, media was replaced with RPMI1640 with 2% v/v B27 with insulin (B27+, Life Technologies 17504044) until Day 16 flow cytometry collection to validate high CM purity. Parallel wells of high purity hPSC-CM differentiations were matured through long-term culture by replacing B27+ media every 2-3 days.

### 4.3 Multi-omic Sample Collection Design

Sample collection was performed on Days 30, 60, and 113 prior to media change. Each sample comprised 1 well of a 12-well plate, which is roughly 1-2 million cells. For RNA, a 1 min incubation with 500 µL cold Trizol reagent (ThermoFisher 15596018) was performed after a DPBS rinse. The Trizol-cell mixture was scraped into 1.5 mL microcentrifuge tubes, snap-frozen in liquid nitrogen, and stored at −80 °C (RNA). Cold (4° C) 100% methanol was used for simultaneous extraction of intracellular metabolites and proteins from a parallel set of wells (parallel to RNA wells)^42^. The proteomic and metabolomic datasets included longitudinal time course maturation samples collected on Days 30, 60, and 113. Additionally, the proteomic and metabolomic datasets included cross-sectional maturation samples collected on Days 99, 117, 133, and 190 from independent differentiations for each cross-sectional time point. These cross-sectional samples were included during data processing workflows, including feature identification, normalization, and other preprocessing steps as indicated below. Only the Day 30, Day 60, and Day 113 samples were included in the downstream multi-omic analyses presented in this study. These three time points represent a defined maturation time course and are supported by corresponding Day 16 flow cytometry data demonstrating high purity differentiation. The remaining time points (Days 99, 117, 133, and 190) originated from independent differentiations with widespread spontaneous contraction but without corresponding flow cytometry characterization and therefore were not included in the maturation time-course analyses described in this manuscript.

### 4.4 Intracellular Metabolite and Protein Extraction

Intracellular metabolites and proteins were collected from a single well using a methanol quench as described previously^42^. Prior to quenching, wells were rinsed twice with DPBS (calcium/magnesium free). To quench metabolic activity, cells were treated for 1 min with 1 mL of cold (4 °C) 100% methanol, and the methanol-cell mixture was scraped into microcentrifuge tubes, vortexed for 10 s, and mixed at 4 °C for 10 min. To create a metabolite supernatant and protein cell pellet, the methanol-cell mixture was centrifuged for 5 min at 14,000g. The protein cell pellet was washed with 200 µL of DPBS. Intracellular metabolite and protein samples were vacuum-dried and stored at −80 °C.

### 4.5 Untargeted Mass Spectrometry-based Intracellular Metabolomics

For sample preparation, intracellular metabolite extracts were thawed on ice and rehydrated with cold 50% methanol prior to mass spectrometry analysis. Reconstitution was performed stepwise by adding 100 µL of 100% methanol with vortexing, followed by 100 µL of mass spectrometry-grade water with vortexing. For data acquisition, flow-injection electrospray Fourier transform ion cyclotron resonance (FIE-FTICR) mass spectrometry-based metabolomics was performed using a Waters ACQUITY UPLC M-Class System (Waters Corporation) coupled to a Bruker solariX 12T FTICR mass spectrometer (Bruker Daltonics) as previously described^139^. Samples were introduced by electrospray ionization. Mass spectrometry-grade solvents were used to prepare mobile phases for positive mode (0.1% formic acid in 50% methanol in water) and negative mode (10 mM ammonium acetate in 50% methanol in water). A 5-minute isocratic flow of 100% pre-mixed mobile phase was delivered at 15 µL/min with a 5 µL injection volume. Fifty scans were collected per spectrum with an ion accumulation time of 0.1 s per scan, resulting in 3.5 minutes of data acquisition per sample, followed by a 1.5-minute wash step. The instrument was calibrated in each ionization mode before analysis using 1 mM NaTFA.

For metabolite detection, data were processed in MetaboScape 2021b (Bruker Daltonics). Bucket lists were generated separately for positive and negative ionization modes using the T-Rex 2D algorithm, then merged using a mass error tolerance of <1.0 ppm. Background signals were removed by excluding features with intensities less than three times those observed in blank samples. SmartFormula annotations were performed in MetaboScape^139^. Features were annotated by accurate mass matching against the Mass Bank of North America and the Human Metabolome Database^52^ using a mass error tolerance of ≤3 ppm. KEGG, HMDB, and PubChem identifiers were assigned to annotated metabolites. All annotated features were manually reviewed in DataAnalysis software (Bruker Daltonics) to confirm feature detection. Features with intensities below 8 × 10^6^ were excluded prior to statistical analysis. At this point, the dataset was restricted to the Day 30, 60, and 113 longitudinal time course maturation samples. Raw metabolite intensity data were processed in R v4.5.2. Missing values were imputed using the quantile regression imputation of left-censored data (QRILC) approach^140^ implemented in the imputeLCMD package v2.1. Intensities were log_2_-transformed^141^ and median-normalized^142^ across samples prior to statistical analysis. Differential metabolite abundance was performed using limma v3.66.0^48^ with multiple comparison p-value adjustment performed using the Benjamini-Hochberg (BH) false discovery (FDR) method. Differentially abundant metabolites were defined by log₂ fold change>1 and padj<0.05. Pathway enrichment analyses were performed using MetaboAnalystR v4.0^51^ or MetaboAnalyst v6.0^50^ and HMDB IDs.

### 4.6 Global Mass Spectrometry-based Quantitative Proteomics

For sample preparation, the dried cell pellet was solubilized in 10 μL of 0.25% w/v Azo^143^, sonicated, subjected to freeze-thaw cycles, and diluted to 0.05% w/v Azo. Protein concentrations were normalized using the Bradford assay. Samples were reduced (10 mM DTT, 1 h, 37 °C) and alkylated (20 mM IAA, 30 min, room temperature in dark), followed by digestion with Trypsin Gold (Promega) at a 50:1 protein-to-enzyme ratio for over 3 h at 37 °C. Enzymatic activity was quenched with formic acid, and Azo was degraded with 305 nm irradiation for 5 min. Peptides were desalted with C18 ZipTips, vacuum-dried, reconstituted in 0.1% formic acid to 0.2 mg/mL, and quantified (NanoDrop - ThermoFisher ND2000c) prior to LC–MS analysis.

For data acquisition, tryptic peptide were analysed on a timsTOF Pro mass spectrometer (Bruker Daltonics) coupled to a nanoElute UHPLC system equipped with a Captivespray nano-electrospray ion source. Approximately 200 ng of peptide was loaded onto an IonOpticks C18 column (25 cm×75 μm i.d., 1.7μm particles) and separated at 55 °C with 400 nL/min flow using a stepwise gradient: 2–17% solvent B (60 min), 17–25% B (60–90 min), and a 30 min wash at 85% B. Solvent A was 0.1% formic acid in water, and solvent B was 0.1% formic acid in acetonitrile. MS and MS/MS data were acquired in Parallel Accumulation–Serial Fragmentation (PASEF) mode^144^ over m/z 100–1700 with ion mobility from 0.60 to 1.60 V·s/cm² (100 ms ramp). Data-dependent acquisition (DDA) included 10 PASEF MS/MS scans per cycle (100% duty cycle). Precursors exceeding 20,000 intensity units were excluded for 0.4 min and reconsidered after 4 min. Collision-induced dissociation (CID) energies were adjusted according to ion mobility (1/K₀). Peptide loading and injection consistency were verified by total ion current comparison to a 200 ng human K562 lysate standard (Promega, 0.2 µg/µL),

Proteins were identified in MaxQuant v1.6.17.0 using the UniProt human database with a 1% FDR at the protein and peptide levels^145^. TIMS-DDA settings included half-width of 6, resolution of 32,000, and precursor mass tolerance ≤30 ppm. Peptides required ≥7 amino acids with a fixed modification for carbamidomethylation and variable modifications for methionine oxidation and N-terminal acetylation. Enzyme specificity was trypsin/P with up to 2 missed cleavages. Label free quantification (LFQ) used a minimum ratio count of 2 with classic normalization. Match between runs used a 0.7 min retention window and 0.05 V·s/cm ion mobility window. MS/MS spectra were required for LFQ quantification. Proteomics data were processed using ProStar and DAPAR R packages^146^. LFQ intensities were log2-transformed. Proteins missing in at least 2 of 3 or 3 of 4 biological replicates across all groups were removed, along with contaminants and reverse hits. Missing values were imputed using the slsa algorithm for partially observed proteins and the 2.5% intensity quantile for proteins absent in one condition. At this point, the dataset was restricted to the Day 30, 60, and 113 longitudinal time course maturation samples. Differential protein expression was performed using limma v3.66.0^48^ with multiple comparison p-value adjustment performed using the Benjamini-Hochberg (BH) false discovery (FDR) method. Differentially expressed proteins were defined by log₂ fold change>1 and padj<0.05. The proteomic CM maturation score was calculated from log_2_-transformed LFQ intensities where a delta between the sample of interest and the mean of Day 30 samples was calculated for each protein displayed in the heatmap. The curated protein set consisted of 113 features of which 68 were detected using proteomics (**Table S1**). Each protein was assigned a directional weight based on its expected change during maturation and the signed deltas were capped at ±1 to prevent outlier influence. The global proteomic CM maturation score for each sample was calculated as the sum of signed, capped deltas across all detected proteins for the protein set displayed in the heatmap. Maturation axis-level and function-level normalized maturation scores were calculated in the same manner for each category of proteins and normalized to the number of detected proteins within the category to account for differences in the number of proteins in each domain. Pathway enrichment analyses were performed using ClusterProfiler v4.14.3^60^. Transcription factor (TF) activity inference was performed using DoRothEA v1.18.0^61^ implemented in decoupleR using all regulons (levels A-E). Univariate linear models between the limma moderated t-statistic and TF regulon weights were used to calculate p-values, which were adjusted using the Benjamini–Hochberg method.

### 4.7 RNA Extraction, Sequencing, and Analysis

RNA was extracted from Trizol samples by phase separation and purified using Direct-zol MiniPrep Plus columns (Zymo R2072) with on-column DNAse treatment. Concentration and purity were assessed with a NanoDrop (A260/A280 ∼2.0; A260/A230 >2.0) and samples were stored at −80°C. Prior to library construction and sequencing, RNA was quantified using Qubit on an Agilent 2100 Bioanalyzer. Libraries were prepared from 500 ng total RNA per sample using the SMARTer Stranded Total RNA Kit - HI Mammalian (Takara) and sequenced on an Illumina NovaSeq 6000. FASTQ files were aligned to the human genome (hg38+decoy) using STAR v2.5.3a^147^. Gene-level counts were generated with featureCounts v2.0.3^148^. Differential expression analysis was performed in R v4.5.2 using DESeq2^49^. Differentially expressed genes were defined by log₂ fold change>1 and padj<0.05. The transcriptomic CM maturation score was calculated from variance-stabilized (VST) expression values where a delta between the sample of interest and the mean of Day 30 samples was calculated for each gene displayed in the heatmap. The curated gene set consisted of 113 features of which all were detected using transcriptomics (**Table S1**). Each gene was assigned a directional weight based on its expected change during maturation and the signed deltas were capped at ±1 to prevent outlier influence. The global transcriptomic CM maturation score for each sample was calculated as the sum of signed, capped deltas across all detected genes for the gene set displayed in the heatmap. Maturation axis-level and function-level normalized maturation scores were calculated in the same manner for each category of genes and normalized to the number of detected genes within the category to account for differences in the number of genes in each domain. Pathway enrichment analyses were performed using ClusterProfiler v4.14.3^60^. Transcription factor (TF) activity inference was performed using DoRothEA v1.18.0^61^ implemented in decoupleR using either high-confidence regulons (levels A and B) or all regulons (levels A-E) as indicated. Univariate linear models between the DESeq2 Wald statistic and TF regulon weights were used to calculate p-values, which were adjusted using the Benjamini–Hochberg method. CIBERSORTx digital cytometry analysis was performed using the Grancharova *et al.* single cell RNA sequencing dataset as a reference using the author-defined cell clusters after removing iPSCs, merging Day 12 and 24 hPSC-CMs into an Early CM cluster, and renaming D90 hPSC-CMs as Late CMs^28,46^.

### 4.8 Integrated Multi-omic Analysis

Multi-omic factor analysis was performed with MOFA2 v1.20.1 using the MEFISTO^83^ framework and only the top 5000 transcripts. Data transformation for each -omic layer included variance stabilizing transformation (VST from DESeq2) for RNA and log2-transformation for metabolomics and proteomics. One-sided (negative or positive factor values) individual -ome (proteomic and transcriptomic) enrichment analyses were performed using MOFA2 and a custom script to convert msigdbr v25.1.1^149^ pathway data frames to binary identity matrices. Multi-omic TF activity inference and comparison to transcriptomic and proteomic TF activity inference was performed using DoRothEA v1.18.0^61^ implemented in decoupleR using all regulons (levels A-E) for all -omes. Integrated Factor 1 weights were calculated by summing transcriptome and proteome weights. Univariate linear models between integrated Factor 1 weights and TF regulon weights were used to calculate p-values, which were adjusted using the Benjamini–Hochberg method. For dual transcriptome and proteome enrichment analyses, parity plots were used to identify genes and proteins with shared expression patterns prior to using ClusterProfiler v4.14.3^60^. For tri-ome analyses using all datasets simultaneously, multi-omic gene set enrichment analysis, multiGSEA v1.16.0^109^ was used.

### 4.9 Flow Cytometry

Cardiomyocyte purity was assessed on D16 via flow cytometry for cardiac troponin T (cTnT) as previously described^7^. Cells were dissociated with Accutase, fixed with 1% paraformaldehyde, and stored in 90% methanol at -20 °C. After washing in flow buffer, (FB = 0.5% w/v BSA in DPBS), cells were incubated overnight at 4°C with msIgG1-anti-cTnT (1:1000, ThermoFisher MA5-12960) in FB + 0.1% v/v Triton X-100. The next day, cells were washed and incubated for 60 min at room temperature in the dark with AF488-anti-msIgG1 (1:1000, ThermoFisher A-21121) in FB + 0.1% v/v Triton X-100. Cells were rinsed and resuspended in 200 µL FB for analysis on a BD Accuri C6 Plus flow cytometer. Undifferentiated hPSC negative control samples were stained in parallel, and gating was performed using FlowJo software.

### 4.10 Immunocytochemistry and Image Analysis

Cells were rinsed with DPBS and fixed for 30 minutes with 1% (v/v) paraformaldehyde at room temperature. Another DPBS rinse was performed prior to blocking with 1% (w/v) BSA in DPBS and storage at 4 °C. After removal from storage, wells were rinsed with DPBS and incubated with primary antibodies overnight in the dark at 4 °C in FB2. The primary antibody solution was rinsed with DPBS, and wells were incubated with secondary antibodies for 60 minutes at room temperature in the dark in FB2. The secondary antibody solution was rinsed with DPBS, and wells were incubated with 1:5000 (2µg/ml) Hoechst 33342 (Invitrogen H3570) in DPBS for 5 minutes. Wells were rinsed with DPBS twice and filled with DPBS for imaging on a Nikon Ti2 microscope equipped with an Aura III light engine (Lumencor 80-10306).

To quantify myosin heavy chain isoforms, we stained with rb-IgG-anti-MYH7 (1:100, R&D Systems MAB90961100) and ms-IgG1-anti-MYH6 (1:100, R&D Systems MAB8979). Primary antibodies were detected with AF488-anti-rbIgG (1:1000, ThermoFisher A11008) and AF647-anti-msIgG1 (1:1000, ThermoFisher A21240). At 20x magnification, the total cardiomyocyte area within MYH7 and MYH6 positive regions was defined using a binary threshold and quantified using Nikon Imaging Software (NIS Elements v5.30.06). Hoechst positive cardiomyocyte nuclei were identified within the MYH7 and MYH6 positive regions using a binary threshold. Nuclear area within double-positive (MYH7+/MYH6+) and single-positive (MYH7+ only or MYH6+ only) regions was used to calculate the percentage of cardiomyocytes in each population (n=8 images per condition). The same procedure was performed to quantify myosin light chain isoforms with different antibody combinations. For myosin light chain isoforms, we stained with ms-IgG2b-anti-MYL7 (1:200, Synaptic Systems 311011, Clone 56F5) and rb-IgG-anti-MYL2 (1:200, Proteintech 10906-1-AP). Primary antibodies were detected with AF488-anti-msIgG2b (1:1000, ThermoFisher A21411) and AF647-anti-rbIgG (1:1000, ThermoFisher A31573).

To quantify sarcomere characteristics and Cx43 puncta in cardiomyocytes, we stained with ms-IgG2b-anti-MF20 (1:25, DSHB), ms-IgG1-anti-alpha-actinin (1:1000, Sigma-Aldrich A7811), and rb-IgG-anti-Cx43 (1:500, Cell Signaling Technology 3512). Primary antibodies were detected using AF488-anti-msIgG2b (1:1000, ThermoFisher A21141), AF647-anti-msIgG1 (1:1000, ThermoFisher A21240), and AF594-anti-rbIgG (1:1000, ThermoFisher A11012). Sarcomere lengths were quantified from α-actinin staining at 60x magnification using ImageJ across 5-8 Z lines (or bands) in 10 sarcomeres for each image (n=8 images per condition). Sarcomere lengths and sarcomere organization scores were also automatically quantified using SotaTool with a minimum sarcomere length of 1.0 µm, a maximum sarcomere length of 2.5µm, 4×4 segmentation, an offset of 4 µm, and default values for all other parameters^47^. For Cx43 puncta in MF20+ cardiomyocytes, imaging at 20x magnification and 1024 resolution was performed on a Nikon A1R+ confocal microscope for a uniform optical section width across images. After converting multiple z-stack images into a maximum intensity projection, cardiomyocyte (MF20), Cx43, and nuclear (Hoechst) channels were converted to binary masks. Masks were intersected to restrict Cx43 quantification to cardiomyocyte regions and to exclude nuclear signal. Particle analysis (size 1-30) was performed in ImageJ within the previously defined intersection of interest for each image (n=4 images per condition, 220 cardiomyocyte nuclei per image).

### 4.11 Mitochondrial/Nuclear DNA Content Analysis

Genomic DNA was isolated from Trizol samples by phase separation. Mitochondrial (mito) and nuclear (nuc) genomic DNA content were quantified by qPCR as previously described^57^. Mitochondrial DNA content was normalized to nuclear DNA, and resulting ratios were normalized to the lowest day control for that differentiation. Primer sequences are as follows for nuclear DNA: Forward – CAACTTCATCCACGTTCACC and Reverse – GAAGAGCCAAGGACAGGTAC. Primer sequences are as follows for mitochondrial DNA: Forward – CGAAAGGACAAGAGAAATAAGG and Reverse - CTGTAAAGTTTTAAGTTTTATGCG

### 4.12 Statistical Analysis

Multi-omics data were collected from 3–4 wells within a WTC11 hiPSC-CM differentiation at each time point. Statistical tests are provided in figure legends. Statistical software for -omic analysis is indicated in the corresponding methods section. Statistical analysis for heatmap-based CM maturation scoring was performed using rstatix v0.7.2. All other statistical comparisons were performed in GraphPad Prism v9.5.1. Annotations include: ∗ (p<0.05), ∗∗ (p<0.01), ∗∗∗ (p<0.001), ∗∗∗∗ (p<0.0001), and ns (p>0.05).

## Data Availability

All data needed to evaluate the conclusions in the paper are present in the paper and/or the Supplementary Materials. Next-generation RNA sequencing data have been deposited on the NCBI Gene Expression Omnibus^150^ under the accession number GSE336395. The mass spectrometry proteomics data have been deposited to the ProteomeXchange Consortium via the PRIDE^151^ partner repository with the dataset identifier PXD079749. The intracellular mass spectrometry metabolomics data have been deposited to Zenodo under the following DOI: doi.org/10.5281/zenodo.20679872.

## Acknowledgements

The authors also would like to acknowledge the UW-Madison Human Proteomics Program Mass Spectrometry Facility (initially funded by the Wisconsin partnership funds) in obtaining mass spectrometry data and NIH S10OD018475 for the acquisition of ultra-high resolution mass spectrometer for biomedical research.

## Author Contributions

Conceptualization: AKF, ADS, SPP

Data Curation: AKF, ADS, EFB, MRP

Formal Analysis: AKF, ADS, EFB, MRP, CP, CJP, FS, XZ

Funding Acquisition: TJK, YG, SPP

Investigation: AKF, ADS, EFB, MRP, YZ, CP, CJP

Methodology: AKF, ADS, EFB, MRP, YZ, CP, CJP, FS, XZ, YG, SPP

Project Administration: JZ, TJK, YG, SPP

Resources: YG, SPP

Software: AKF, ADS Supervision: JZ, TJK, YG, SPP

Validation: AKF, ADS, EFB, YG, SPP

Writing—original draft: AKF

Writing—review & editing: All authors

