## Supplemental Figures for "Multi-omic analysis reveals maturation programs in human pluripotent stem cell-derived cardiomyocytes during long-term culture"

### Supplemental Material

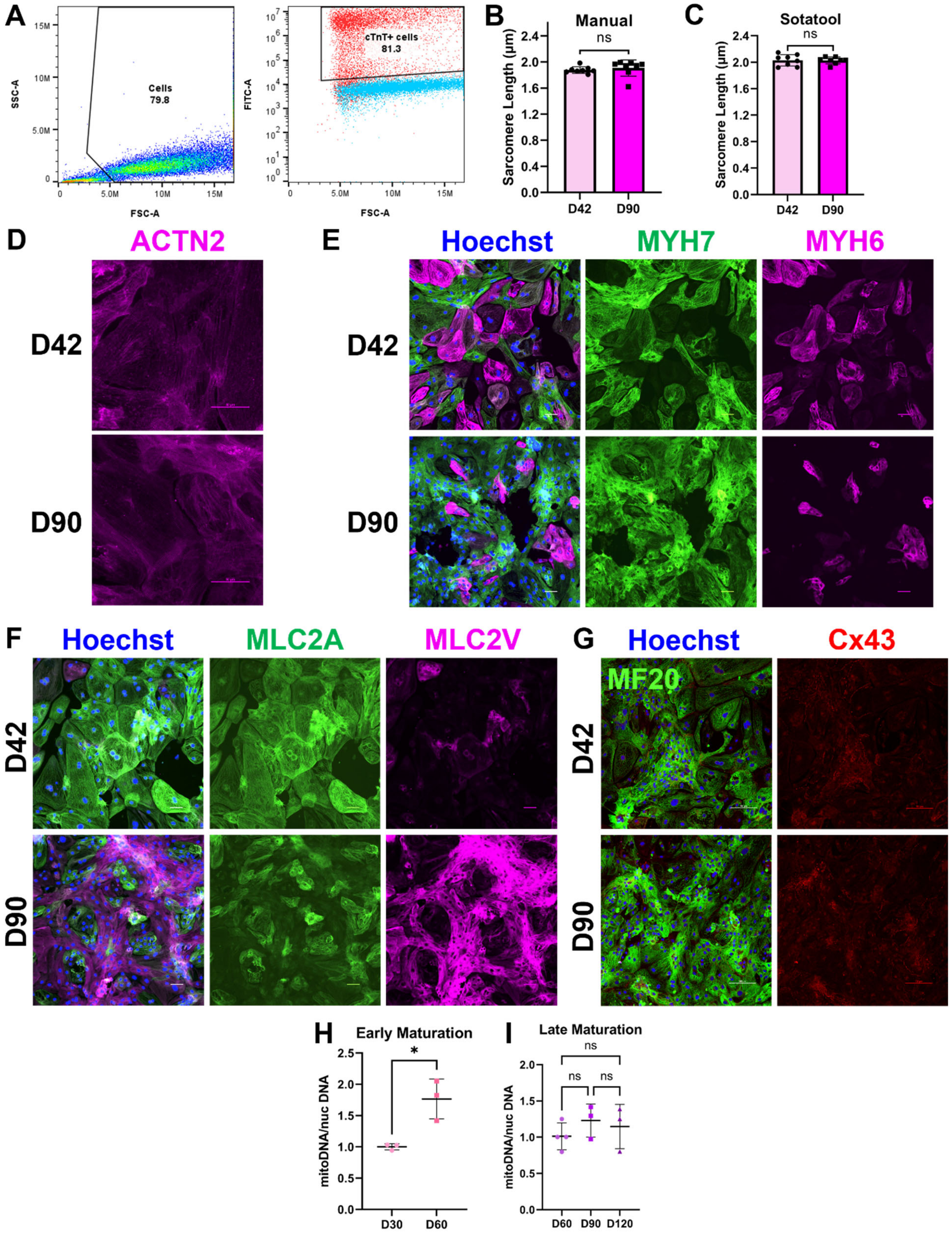

**Figure S1. Flow cytometry gating and hPSC-CM maturation phenotypes during long-term culture.** **A)** Representative cell gating (SSC-A vs. FSC-A). Representative cTnT gating (FITC-A vs. FSC-A) with hPSC-CM sample in red and hPSC negative control in blue. **B)** Sarcomere length quantified from ACTN2 images using manual measurement. **C)** Sarcomere length quantified from ACTN2 images using automated measurement with SotaTool. N=8 images per time point. P-values from unpaired t-tests. **D)** Representative ACTN2 sarcomere images at 60x magnification. Scale bars = 50 µm. **E)** Representative MYH7/MYH6 isoform images at 20x magnification. Scale bars = 50 µm. **F)** Representative MLC2V/MLC2A isoform images at 20x magnification. Scale bars = 50 µm. **G)** Representative MF20 and Cx43 images at 20x magnification. Scale bars = 100 µm. **H)** Mitochondrial-to-nuclear DNA content normalized to D30 ($2^{-(CT_{D60}-CT_{D30})}$) for an early maturation trajectory. N=3 replicates per time point. P-value from unpaired t-test. **I)** Mitochondrial-to-nuclear DNA content normalized to D60 ($2^{-(CT_{D120\text{ or }D90}-CT_{D60})}$) for a late maturation trajectory. N=3-4 replicates per time point. P-values from one-way ANOVA with Tukey’s post-hoc tests.

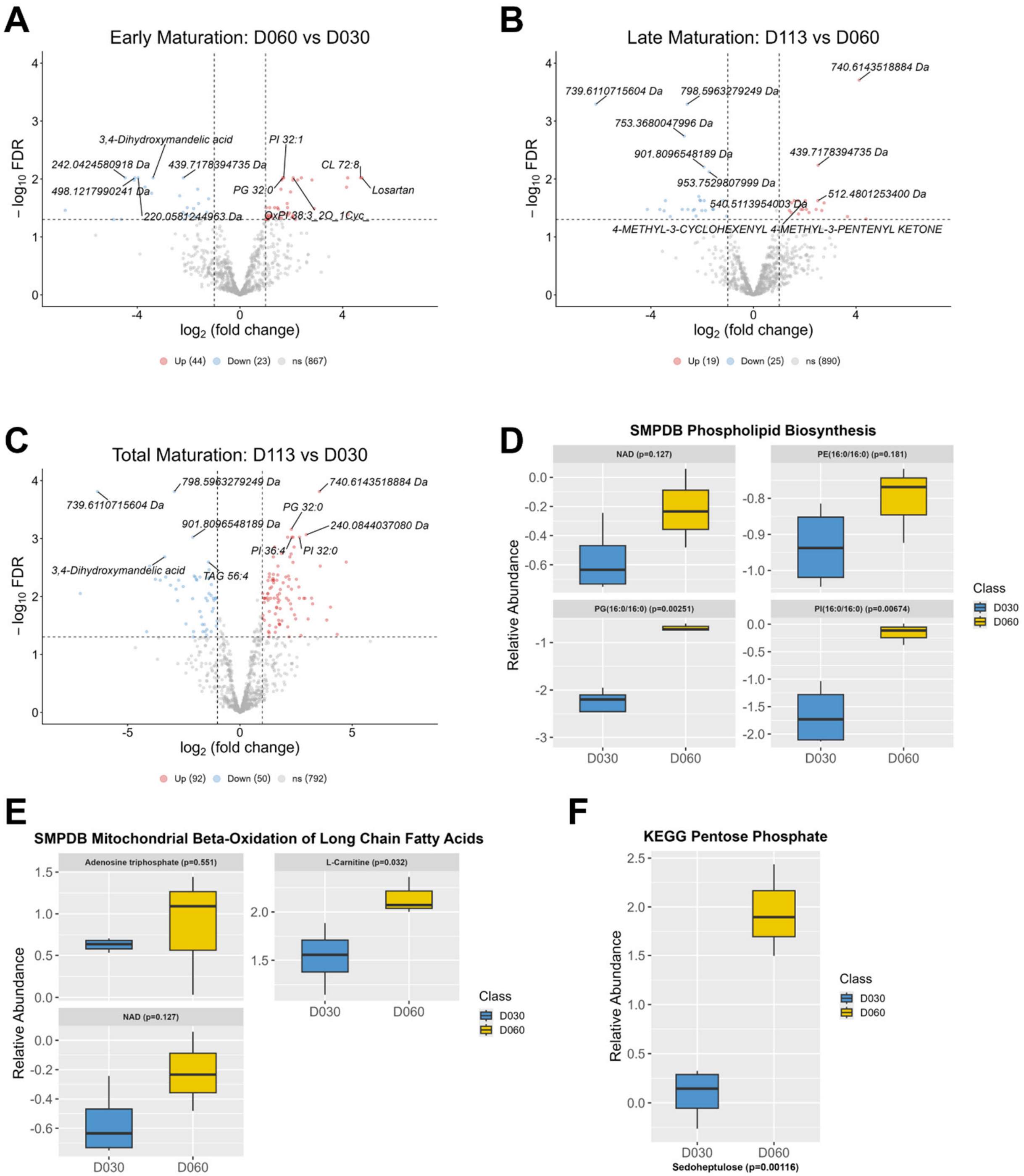


**Figure S2. Differentially abundant metabolites and Day 60 vs. Day 30 early maturation pathway enrichment.**

**A)** Volcano plot for metabolomic early maturation comparison Day 60 vs. Day 30. **B)** Volcano plot for metabolomic late maturation comparison Day 113 vs. Day 60. **C)** Volcano plot for metabolomic total maturation comparison Day 113 vs. Day 30. Dotted lines for all volcano plots represent significance thresholds of absolute value of $\log_2 FC > 1$ and FDR-adjusted p-value $< 0.05$. Metabolites in red are upregulated (Up) in the later time point. Metabolites in blue are downregulated (Down) in the later time point. Metabolites in gray are not significantly changed (ns). **D)** Relative abundances of metabolites in the SMPDB Phospholipid Biosynthesis pathway, which was significant for the quantitative enrichment analysis between D60 and D30. **E)** Relative abundance of metabolites in the SMPDB Mitochondrial Beta-Oxidation of Long Chain Fatty Acids pathway, which was significant for quantitative enrichment between D60 and D30. **E)** Relative abundance of sedoheptulose in the KEGG Pentose Phosphate pathway, which was significant for quantitative enrichment between D60 and D30. All box plots display log2-transformed, normalized metabolite levels for each metabolite at D30 and D60 (N=3-4 replicates per time point) and p-values are from unpaired t-tests.

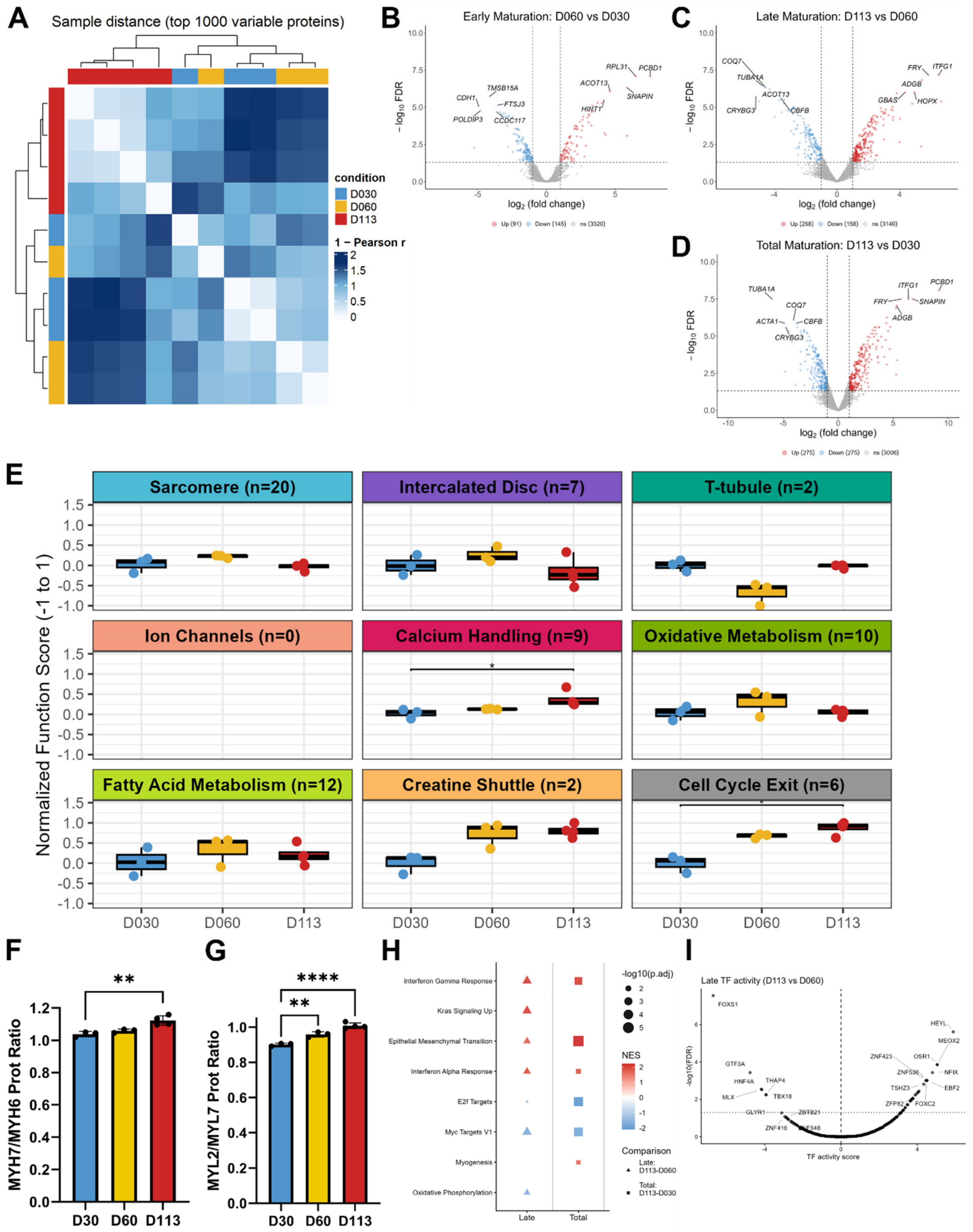


**Figure S3. Differentially abundant proteins, function-level maturation scores, pathway enrichment, and TF regulon analyses.**

A) Heatmap of sample distances based on the top 1000 most variable proteins calculated using Pearson correlation with lighter colors indicating higher pairwise similarity between samples. **B)** Volcano plot for proteomic early maturation comparison Day 60 vs. Day 30. **C)** Volcano plot for proteomic late maturation comparison Day 113 vs. Day 60. **D)** Volcano plot for proteomic total maturation comparison Day 113 vs. Day 30. Dotted lines for all volcano plots represent significance thresholds of absolute value of log2FC > 1 and FDR-adjusted p-value < 0.05. Proteins in red are upregulated (Up) in the later time point. Proteins in blue are downregulated (Down) in the later time point. Proteins in gray are not significantly changed (ns). **E)** Normalized function-level maturation scores across time in culture. Scores were calculated from signed, capped sample deltas in log2 LFQ intensities and each axis score was normalized by the number of proteins. P-values from Kruskal Wallis with a Dunn's post hoc test. **F)** MYH7/MYH6 ratio of protein expression from $log_2$-transformed LFQ intensities. **G)** MYL2/MYL7 ratio of protein expression from $log_2$-transformed LFQ intensities. N=3-4 replicates per time point. Sarcomere isoform ratio p-values from one-way ANOVA with Dunnett's post hoc test. **H)** Hallmark GSEA of proteomic changes. Dot size indicates -log10(padj) and color indicates normalized enrichment score. **I)** DoRothEA TF activity volcano plot for proteomic late maturation comparison Day 113 vs. Day 60. TF activity was inferred using univariate linear modeling (ULM) with Benjamini-Hochberg FDR-adjusted p-values and included all regulons (levels A-E).

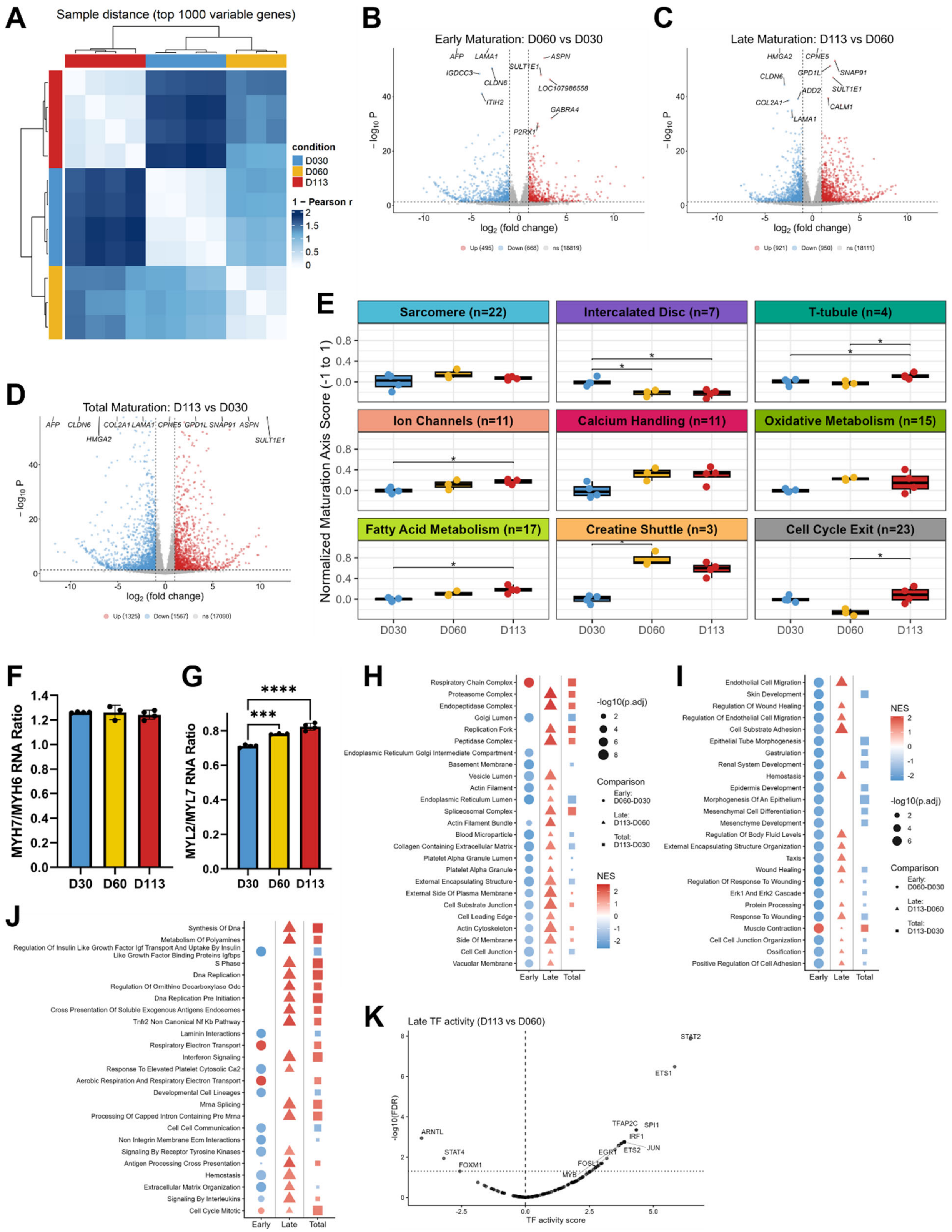

**Figure S4. Differentially abundant transcripts, function-level maturation scores, pathway enrichment, and TF regulon analyses.**

**A)** Heatmap of sample distances based on the top 1000 most variable transcripts calculated using Pearson correlation with lighter colors indicating higher pairwise similarity between samples. **B)** Volcano plot for transcriptomic early maturation comparison Day 60 vs. Day 30. **C)** Volcano plot for transcriptomic late maturation comparison Day 113 vs. Day 60. **D)** Volcano plot for transcriptomic total maturation comparison Day 113 vs. Day 30. Dotted lines for all volcano plots represent significance thresholds of absolute value of log2FC > 1 and FDR-adjusted p-value < 0.05. Transcripts in red are upregulated (Up) in the later time point. Transcripts in blue are downregulated (Down) in the later time point. Transcripts in gray are not significantly changed (ns). **E)** Normalized function-level maturation scores across time in culture. Scores were calculated from signed, capped sample deltas in VST-transformed counts and each axis score was normalized by the number of transcripts. P-values from Kruskal Wallis with a Dunn's post hoc test. **F)** *MYH7/MYH6* ratio of gene expression from VST-transformed counts. **G)** *MYL2/MYL7* ratio of gene expression from VST-transformed counts. N=3-4 replicates per time point. Sarcomere isoform ratio p-values from one-way ANOVA with Dunnett's post hoc test. **H)** GO Cellular Component GSEA of transcriptomic changes. **I)** GO Biological Process GSEA of transcriptomic changes. **J)** Reactome GSEA of transcriptomic changes. Dot size indicates -log10(padj) and color indicates normalized enrichment score. **K)** DoRothEA TF activity volcano plot for transcriptomic late maturation comparison Day 113 vs. Day 60. TF activity was inferred using univariate linear modeling (ULM) with Benjamini-Hochberg FDR-adjusted p-values and restricted to high-confidence regulons (levels A and B).

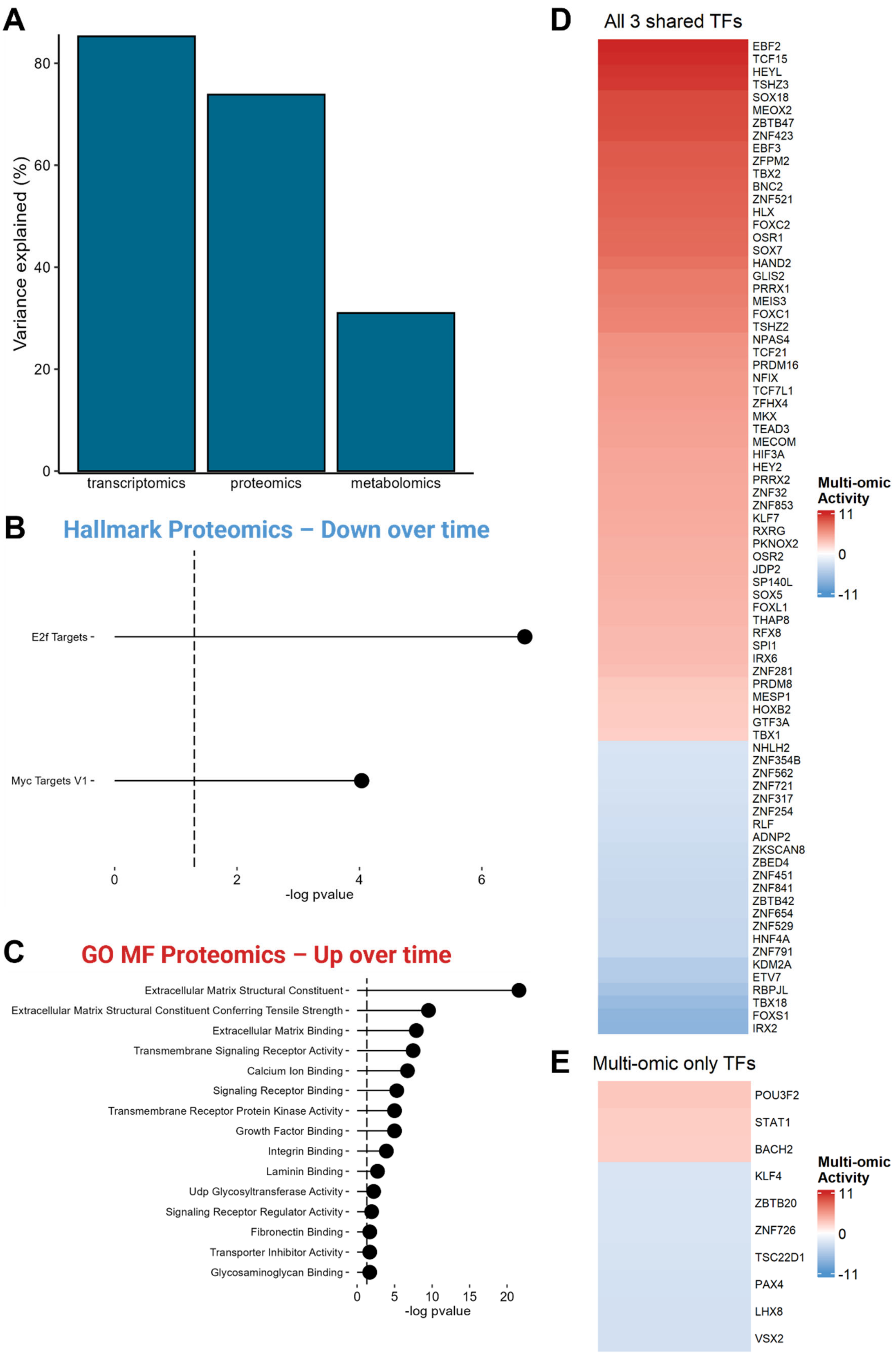
A
Variance explained (%)
0
20
40
60
80
transcriptomics
proteomics
metabolomics
B
Hallmark Proteomics – Down over time
E2f Targets
Myc Targets V1
0
2
4
6
-log pvalue
C
GO MF Proteomics – Up over time
Extracellular Matrix Structural Constituent
Extracellular Matrix Structural Constituent Conferring Tensile Strength
Extracellular Matrix Binding
Transmembrane Signaling Receptor Activity
Calcium Ion Binding
Signaling Receptor Binding
Transmembrane Receptor Protein Kinase Activity
Growth Factor Binding
Integrin Binding
Laminin Binding
Udp Glycosyltransferase Activity
Signaling Receptor Regulator Activity
Fibronectin Binding
Transporter Inhibitor Activity
Glycosaminoglycan Binding
0
5
10
15
20
-log pvalue
D
All 3 shared TFs
EBF2
TCF15
HEYL
TSHZ3
SOX18
MEOX2
ZBTB47
ZNF423
EBF3
ZFPM2
TBX2
BNC2
ZNF521
HLX
FOXC2
OSR1
SOX7
HAND2
GLIS2
PRRX1
MEIS3
FOXC1
TSHZ2
NPAS4
TCF21
PRDM16
NFIX
TCF7L1
ZFHX4
MKX
TEAD3
MECOM
HIF3A
HEY2
PRRX2
ZNF32
ZNF853
KLF7
RXRG
PKNOX2
OSR2
JDP2
SP140L
SOX5
FOXL1
THAP8
RFX8
SPI1
IRX6
ZNF281
PRDM8
MESP1
HOXB2
GTF3A
TBX1
NHLH2
ZNF354B
ZNF562
ZNF721
ZNF317
ZNF254
RLF
ADNP2
ZKSCAN8
ZBED4
ZNF451
ZNF841
ZBTB42
ZNF654
ZNF529
HNF4A
ZNF791
KDM2A
ETV7
RBPJL
TBX18
FOXS1
IRX2
Multi-omic Activity
11
0
-11
E
Multi-omic only TFs
POU3F2
STAT1
BACH2
KLF4
ZBTB20
ZNF726
TSC22D1
PAX4
LHX8
VSX2
Multi-omic Activity
11
0
-11

**Figure S5. MOFA proteomic pathway analysis and multi-omic TF activity over time in culture.**
**A)** Percent variance explained by each -ome across all factors. **B)** Hallmark GSEA for negative proteome factor values. **C)** GO MF GSEA for positive proteome factor values. **D)** Heatmap of DoRothEA multi-omic TF activity for the 78 TFs significantly changed across the proteome, transcriptome and multi-ome. TF analysis for all DoRothEA regulons (levels A-E). **E)** Heatmap of DoRothEA multi-omic TF activity for the 10 TFs significantly changed across multi-ome. DoRothEA TF analysis for all regulons (levels A-E). Multi-omic TF activity is inferred from the summation of MOFA Factor 1 weights (correlated with time in culture) across the proteome and transcriptome. Positive (red) Multi-omic Activity indicates that the TF activity is upregulated with time (Factor 1). Negative (blue) Multi-omic Activity indicates that the TF activity is downregulated with time (Factor 1).

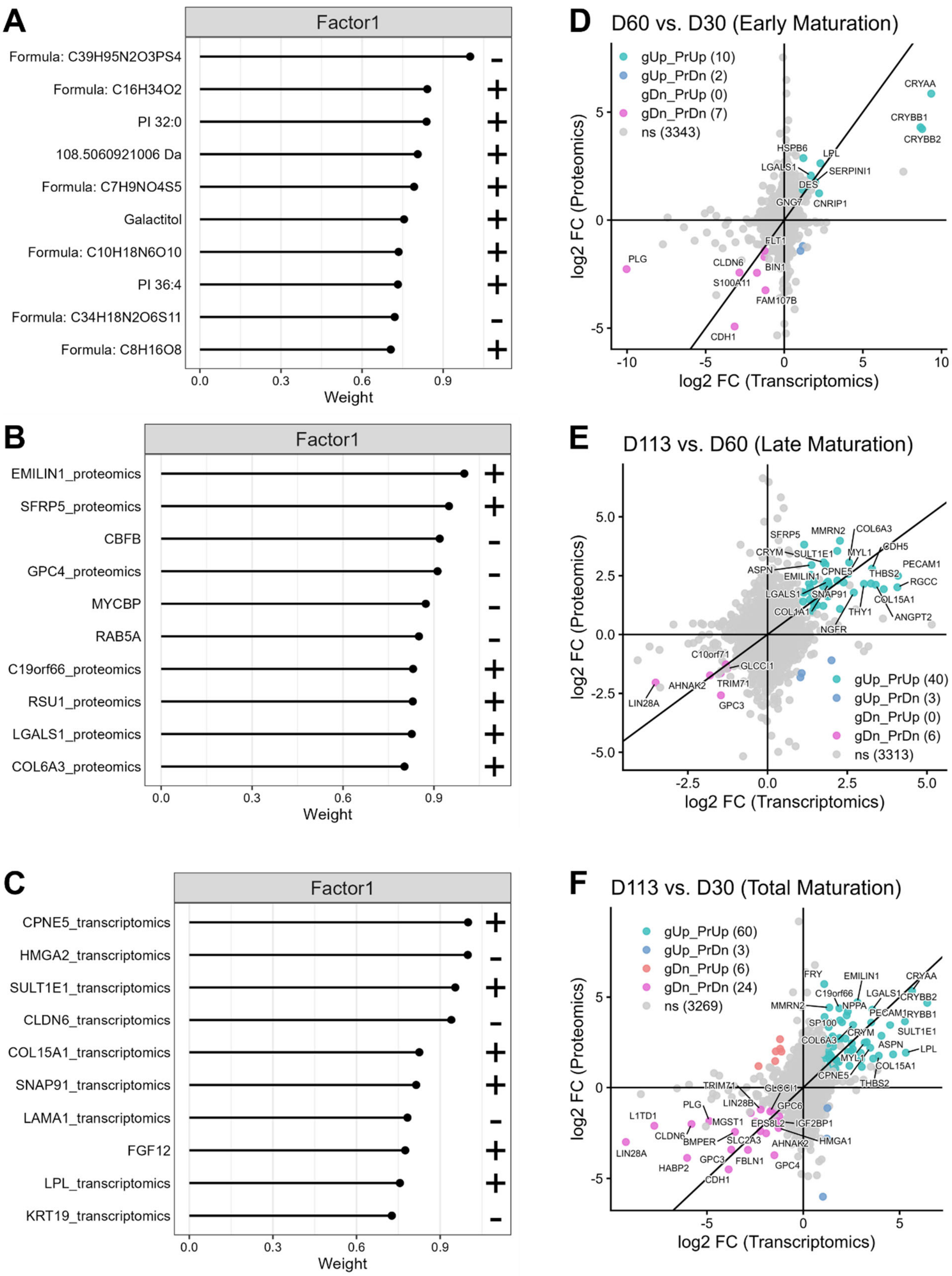


**Figure S6. Multi-omic feature correlation with Factor 1 and concordant proteome and transcriptome changes over time in culture.**

**A)** Top 10 metabolites correlated with Factor 1 values. Sign indicates direction of change with increased Factor 1 value (time). **B)** Top 10 proteins correlated with Factor 1 values. Sign indicates direction of change with increased Factor 1 value (time). **C)** Top 10 transcripts correlated with Factor 1 values. Sign indicates direction of change with increased Factor 1 value (time). **D)** Parity plot comparing proteome and transcriptome $log_2$ fold change (log2 FC) for the D60 vs. D30 early maturation comparison. **E)** Parity plot comparing proteome and transcriptome $log_2$ fold change (log2 FC) for the D113 vs. D60 late maturation comparison. **F)** Parity plot comparing proteome and transcriptome $log_2$ fold change (log2 FC) for the D113 vs. D30 total maturation comparison. A gene-protein pair indicated as significantly up or down required an FDR-adjusted p-value < 0.05 in both the transcriptome and proteome as well as a log2 FC greater than the absolute value of 1. gUp = gene upregulated. PrUp = protein upregulated. gDn = gene downregulated. PrDn = protein downregulated. ns = gene and/or protein changes are not significant.

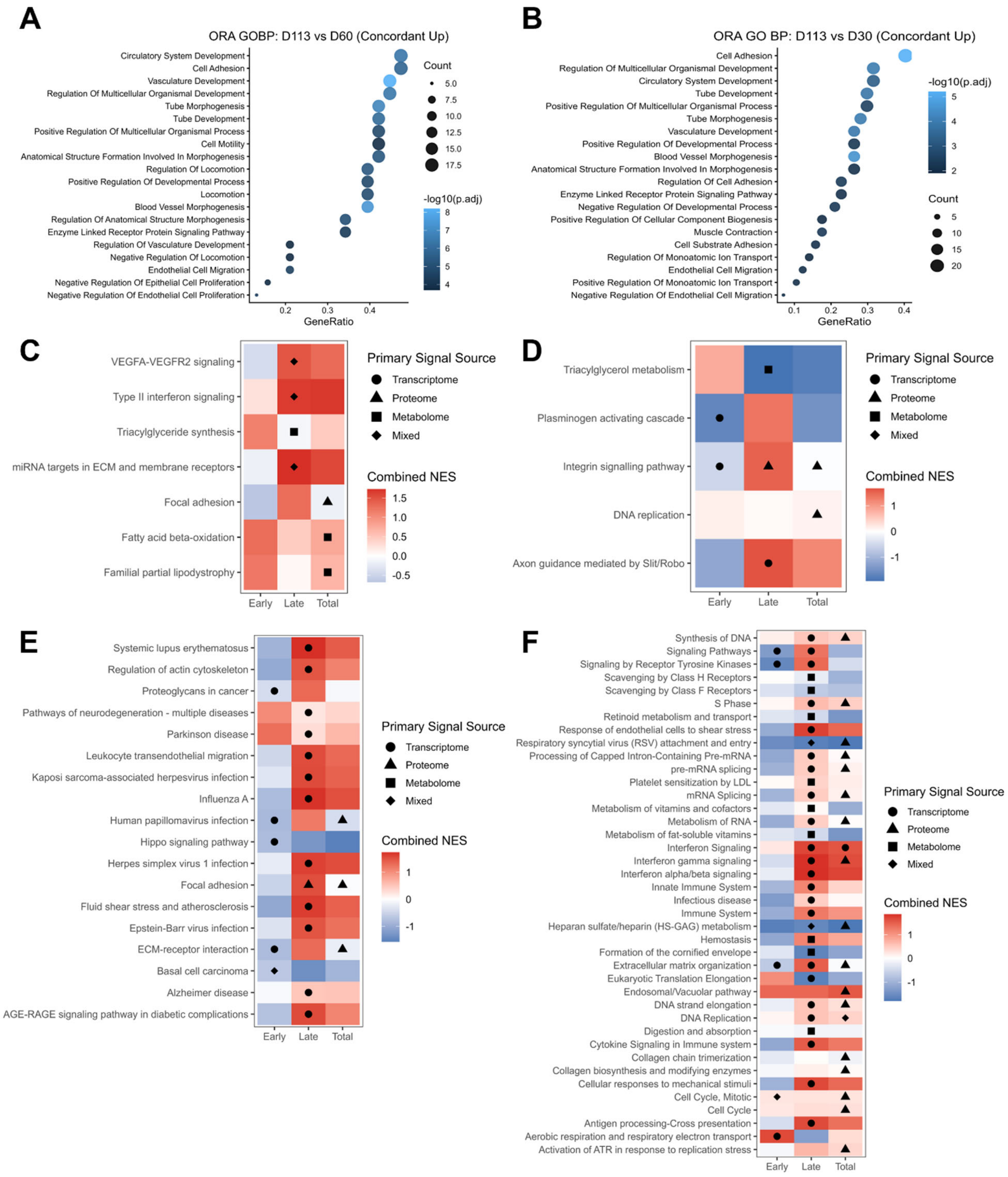


**Figure S7. Pathway analysis of concordant proteome and transcriptome changes and MultiGSEA pathway analysis.**
**A)** GO BP ORA for gene-protein pairs (N=40) concordantly upregulated (gUp_PrUp) in the D113 vs. D60 late maturation comparison. **B)** GO BP ORA for gene-protein pairs (N=60) concordantly upregulated (gUp_PrUp) in the D113 vs. D30 total maturation comparison. **C)** WikiPathways MultiGSEA pathway analysis for all significantly enriched pathways across the metabolome, proteome, and transcriptome. **D)** Panther MultiGSEA pathway analysis for all significantly enriched pathways across the metabolome,

proteome, and transcriptome. **E)** KEGG MultiGSEA pathway analysis for all significantly enriched pathways across the metabolome, proteome, and transcriptome. **F)** Reactome MultiGSEA pathway analysis for the top 40 significantly enriched pathways across the metabolome, proteome, and transcriptome. Combined P-value from Stouffer's Z-score method for p-value combination across -omes. Combined normalized enrichment score (NES) was calculated as the weighted mean of -ome specific NES values, with weight contributions of each -ome equal to the square root of the number of detected pathway features identified in a specific -ome. Primary -omic signal source identified from the lowest -ome specific adjusted p-value if $< 0.10$ and -ome specific NES has absolute value greater than 1. If both conditions are not true, then the -omic signal is labeled as mixed. The number of mapped features in each -ome for MultiGSEA: 150 metabolites, 3374 proteins, and 18009 transcripts.
